# Rewiring of 3D Enhancer-Promoter Interactome Underlies Diabetic Endothelial Dysfunction

**DOI:** 10.64898/2026.03.03.709273

**Authors:** Lei Jiang, Xiaofei Yang, Ruixin Zhou, Shiyu Zheng, Yelan Li, Ning Tan, Siim Pauklin, Sakthivel Sadayappan, Chunxiang Zhang, Wanzi Hong, Mingyang Wang, Hannah Morgan, Keara Little, Guochang Fan, Furong Li, Anil G. Jegga, Jinsong Bian, Gangjian Qin, Wei Huang, Liuyang Cai, Yuliang Feng

**Author notes:** These authors contributed equally to this work (Lei Jiang, Xiaofei Yang, Ruixin Zhou). Correspondence: Lei Jiang, Wei Huang, Liuyang Cai, Yuliang Feng.

## Abstract

**Background:** Diabetic vascular complications are driven by endothelial dysfunction, yet the role of 3D genome organization in this process is unknown. We sought to define the alterations in chromatin architecture in diabetic endothelium and identify the key regulators involved.

**Methods:** We generated a high-resolution 3D epigenomic atlas of diabetic endothelial cells from mouse models and human subjects using H3K27ac HiChIP, complemented by ChIP-seq, ATAC-seq, and RNA-seq. A human cohort was used to assess protein expression in diabetic versus non-diabetic endothelial cells. To identify JUNB-interacting proteins, we performed rapid immunoprecipitation mass spectrometry of endogenous proteins (RIME), with protein-protein interaction validated by co-immunoprecipitation. Functional validation was performed using *in vitro*, *ex vivo*, and *in vivo* approaches, including endothelial-specific knockdown in a diabetic hindlimb ischemia model.

**Results:** Multi-omics profiling revealed extensive enhancer reprogramming in diabetic endothelium, with AP-1 binding motifs being consistently and selectively enriched in downregulated enhancers across three distinct diabetic models. Analysis of a human cohort confirmed significantly reduced JUNB protein levels in diabetic endothelial cells. We identified widespread disruption of JUNB-anchored enhancer-promoter interactions, which underlies transcriptional repression of key endothelial genes. RIME and co-immunoprecipitation established the E3 ubiquitin ligase RBBP6 as a direct JUNB interactor that promotes its polyubiquitination and proteasomal degradation in response to hyperglycemia. Human cohort analysis further showed reciprocal elevation of RBBP6 in diabetic endothelial cells. Either *JUNB* overexpression or *RBBP6* knockdown restored enhancer-promoter connectivity, reactivated vasoprotective transcriptional programs, and rescued endothelial function. Critically, endothelial-specific knockdown of *Rbbp6* in diabetic mice restored endothelium-dependent vasorelaxation and improved perfusion recovery after hindlimb ischemia, independent of systemic glucose levels.

**Conclusions:** Our study unveils a novel mechanism whereby hyperglycemia induces enhancer reprogramming and disrupts endothelial 3D genome architecture through RBBP6-mediated degradation of JUNB. The RBBP6-JUNB axis is established as a crucial link between metabolic stress and epigenomic reprogramming in vascular disease, presenting a promising therapeutic target for diabetic vasculopathy.

## Introduction

Diabetic vascular complications are the leading cause of morbidity and mortality in diabetes^1^. While endothelial dysfunction is recognized as a central driver of these complications, the molecular mechanisms underlying persistent vascular impairment remain incompletely understood^2^. Emerging evidence suggests that metabolic disturbances in diabetes induce lasting epigenetic changes that disrupt endothelial homeostasis^3,4^, yet how these alterations reshape global chromatin architecture remains unexplored.

Transcription factors, such as AP-1 family^5^, have been implicated in maintaining endothelial function. However, its potential role in orchestrating the three-dimensional (3D) chromatin architecture essential for endothelial gene regulation has not been investigated. Recent advances in 3D genomics have revolutionized our understanding of gene regulation by revealing how spatial genome organization enables precise transcriptional control^6–8^. Landmark studies, including work by Su et al.^9^, have demonstrated the power of chromatin interaction maps in elucidating cell-type-specific regulatory programs in metabolic diseases. Nevertheless, a critical gap remains in our knowledge of how diabetes alters endothelial chromatin topology to drive vascular pathology.

Here, we present the first high-resolution 3D genome atlas of diabetic endothelium, generated through integrative multi-omics analyses. Our findings uncover a previously unrecognized mechanism of endothelial dysfunction in diabetes involving RBBP6-mediated JUNB degradation and subsequent chromatin disorganization. We demonstrate that JUNB serves as a critical architectural factor maintaining endothelial enhancer-promoter interactions, and that its depletion in diabetes leads to selective loss of chromatin loops at genes essential for vascular homeostasis. Importantly, we show that perturbation of RBBP6 restores JUNB protein expression, rescuing both chromatin organization and endothelial function. These discoveries provide a novel framework for understanding diabetic vasculopathy, positioning chromatin topology as a key interface between metabolic stress and endothelial dysfunction.

Our work not only establishes 3D genome disorganization as a hallmark of diabetic endothelium but also reveals the RBBP6-JUNB axis as a potential therapeutic target for vascular complications. By linking metabolic dysregulation to chromatin-based mechanisms of gene control, this study opens new avenues for developing targeted interventions to preserve vascular health in diabetes.

## Methods

### Data availability

Raw data from H3K27ac ChIP-seq, JUNB ChIP-seq, RNA-seq and ATAC-seq have been deposited at Gene Expression Omnibus (GEO) database (GEO: GSE291636). All other relevant data supporting the key findings of this study are available within the article and its Supplementary Information files. Any additional information in this paper is available from the lead contact upon request. Source data are provided with this paper. Detailed Methods and Materials are provided in the Supplemental Material. Please see the Major Resources Table in the Supplemental Material.

## Results

### Loss of JUNB-centric promoter-enhancer interactions is a hallmark of diabetic endothelial cells

To delineate the chromatin architecture of endothelial cells in diabetes, we performed comprehensive 3D-epigenomic analysis in aortic endothelial cells from three diabetic mouse models: type I (Streptozotocin-induced C57BL6, STZ) and two type II diabetic mouse models (C57BL6 db/db and BKS db/db), along with their specific control mice (**Figure 1A**). First, we performed H3K27ac (a prominent histone mark for active enhancers and promoters^10–12^) ChIP-seq analysis to explore the landscape of active cis-regulatory elements (CREs). Principal Component Analysis (PCA) for H3K27ac ChIP-seq revealed clear separation between diabetic and control endothelial cells across all three mouse models, confirming consistent and distinct chromatin signatures (**Figure S1A, Table S1**). Through differential peak analysis, we uncovered a widespread reprogramming of cis-regulatory elements (CREs) in diabetic endothelial cells across the BKS db/db (n = 7,633), C57BL6 db/db (n = 5,701), and STZ (n = 4,313) models, underscoring the profound epigenomic alterations associated with diabetic conditions (**Figure 1B**).

**Figure 1.**
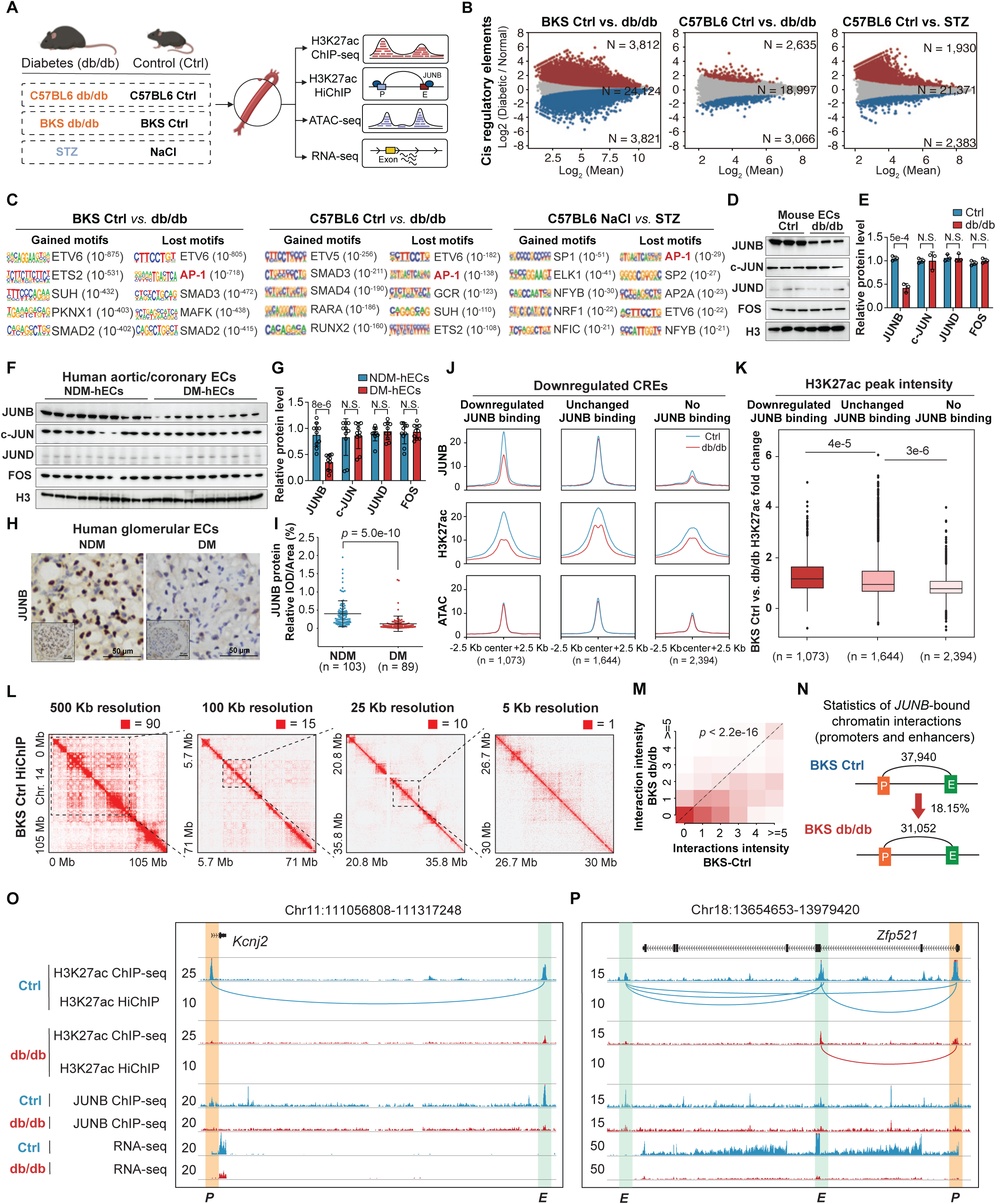
Loss of JUNB-centric promoter-enhancer interactions was a hallmark of diabetic endothelial dysfunction. A. Schematic of experimental design using three diabetic mouse models (type I STZ, type II C57BL6 db/db, and type II BKS db/db) alongside their respective controls). Multi-omics datasets, including H3K27ac HiChIP, H3K27ac ChIP-seq, RNA-seq, and ATAC-seq, were generated. B. MA plots showing differential H3K27ac ChIP-seq peaks of three diabetic mouse models. C. Transcription factor motif enrichment analysis of cis-regulatory elements (CREs) showing the top 5 enriched motifs in both gained and lost enhancers in diabetic mouse endothelial cells. See also Table S2. D. Western blot analysis of AP-1 family members (JUNB, c-JUN, JUND, and FOS) in diabetic mouse aortic endothelial cells (db/db) versus control aortic endothelial cells (Ctrl). E. Quantitative analysis of AP-1 family member expression by Western blotting in endothelial cells from db/db (n = 3) and Ctrl (n = 3) mice. Data were analyzed by Student’s *t* test. F. Western blot analysis of AP-1 family members (JUNB, c-JUN, JUND, and FOS) in human diabetic aortic/coronary endothelial cells (DM-hECs) versus normal human aortic/coronary endothelial cells (Normal-hECs). G. Quantitative analysis of AP-1 family member expression by Western blotting in diabetic (n = 10) and normal (n = 10) human aortic/coronary endothelial cells. Data were analyzed by Student’s *t* test. H. Representative immunohistochemical staining of JUNB in human glomerular endothelial cells from non-diabetic (NDM) and diabetic (DM) patients. I. Quantification of JUNB expression in human glomerular endothelial cells from non-diabetic (NDM, n = 103) and diabetic patients (DM, n = 89). Data were analyzed by Student’s *t* test. J. Comparison of JUNB ChIP-seq, H3K27ac ChIP-seq, and ATAC-seq signal intensities among three groups of downregulated CREs in db/db verus Ctrl mice: downregulated JUNB binding (n = 1,073), unchanged JUNB binding (n = 1,644), and no JUNB binding (n = 2,394). K. Fold change in H3K27ac ChIP-seq signals (BKS Ctrl vs. dbdb) in mouse endothelial cells at CREs in **Figure 1J**. The CREs are grouped by JUNB binding alterations: downregulated (n = 1,073), unchanged (n = 1,644) and no binding (n = 2,394). Data were analyzed by Wilcoxon rank sum test with continuity correction. L. H3K27ac HiChIP interaction maps of BKS Ctrl endothelial cells at 500 kb,100 kb, 25 kb and 5 kb resolutions. Warmer colors (red) indicate higher contact frequencies, and cooler colors (white) represent lower contact frequencies. M. Comparison of chromatin interaction intensity of JUNB-bound CREs in endothelial cells from BKS db/db mice (n = 2) versus Ctrl mice (n = 2). Data were analyzed by Wilcoxon rank sum test with continuity correction. N. Summary of chromatin interaction counts in BSK Ctrl (n = 37,940) and db/db (n = 31,052) HiChIP datasets. O-P. Genome browser view displaying multi-omics datasets including H3K27ac ChIP-seq signals, H3K27ac HiChIP chromatin interactions, JUNB ChIP-seq signals and RNA-seq expression values at the *Kcnj2* (O) and *Zfp521* (P) loci in BKS Ctrl and db/db endothelial cells.

We next examined the underlying transcription factor binding motifs and gained mechanistic insight into the regulatory factors driving these CRE alterations. The motif analysis in both upregulated and downregulated cis-regulatory elements (CREs) across three diabetic models revealed a consistent and significant enrichment of AP-1 binding motifs in downregulated CREs (**Figures 1C**, **Table S2**), indicating that impaired AP-1–mediated transcriptional regulation is a shared epigenetic hallmark of diabetic endothelial dysfunction. The AP-1 motif is a well-characterized binding site for AP-1 family transcription factors, including JUNB, c-JUN, JUND, and FOS^13^. To determine which specific AP-1 factor might drive CRE dysregulation in diabetes, we measured the protein expression level of JUNB, c-JUN, JUND, and FOS across multiple diabetic endothelial cells. Notably, in aortic endothelial cells isolated from BKS db/db mice (db/db), the JUNB protein level was significantly decreased relative to control endothelial cells (Ctrl), whereas the expression of c-JUN, JUND, and FOS remained unchanged (**Figure 1D and 1E**). Consistent with the mouse data, JUNB protein level was also significantly lower in aortic/coronary endothelial cells derived from diabetic patients (DM, n = 10), compared to those in non-diabetic individuals (NDM-hECs, n = 10) (**Figure 1F and 1G**). Furthermore, immunohistochemical analysis of a human cohort revealed significantly lower JUNB expression in glomerular endothelial cells from diabetic patients (DM-hECs, n = 89) compared to non-diabetic individuals (NDM, n = 103) (**Figure 1H and 1I**). Therefore, we performed JUNB ChIP-seq in BKS mice. Interestingly, JUNB ChIP-seq data revealed that JUNB-bound CREs exhibited greater H3K27ac activity than non-JUNB bound CREs (**Figure S1B**). We characterized downregulated CREs into three distinct groups based on JUNB occupancy: downregulated JUNB binding, unchanged JUNB binding, no JUNB binding (**Figure 1J**). Among these groups, the CREs with downregulated JUNB binding showed the greatest reduction in H3K27ac peak intensity compared to the other two groups (**Figure 1K**), highlighting a significant association between the loss of JUNB binding and decreased CRE activity in diabetic endothelial cells, and that JUNB downregulation likely contributes to diabetic endothelial dysfunction.

Given that enhancer-promoter (E-P) interactions are fundamental for precise transcriptional control and cell-type-specific gene expression programs^6^, it is crucial to elucidate how JUNB specifically influences these long-range chromatin interactions for endothelial homeostasis. To address this, we first performed proximity ligation-based global chromatin conformation profiling-HiChIP^14–18^ by mapping H3K27ac-centric chromatin interactions in aortic endothelial cells from BKS Ctrl mice (**Table S1**). The analysis of contact matrices at multiple resolution levels revealed characteristic genome organization features, including compartments at 500-kb resolution, topologically associating domains (TADs) at 100-kb, 25-kb resolution, and focal chromatin loops at 5-kb, 1-kb resolution (**Figure 1L**), consistent with previously published HiChIP findings^19,20^.Gene Ontology (GO) analysis revealed that the genes linked to JUNB-bound anchors were significantly enriched for endothelial-specific functions, including regulation of sprouting angiogenesis and endothelial cell differentiation, compared to those linked to JUNB-unbound anchors (**Figure S1C and S1D, Table S3**), highlighting the crucial role of JUNB-mediated E-P interactions in endothelial gene regulation. Subsequently, we performed HiChIP experiments in aortic endothelial cells from BKS db/db mice, and PCA revealed a clear separation between chromatin loops of diabetic and control mice (**Figure S1E**). Notably, the JUNB-bound E-P interactions exhibited markedly reduced contact strength in BKS db/db endothelial cells compared to controls (**Figure 1M**). Consistent with the observed reduction in interaction intensity, the total number of JUNB-bound E–P loops was also decreased in BKS db/db endothelial cells (n = 31,052) compared to BKS Ctrl cells (n = 37,940) (**Figure 1N**), indicating reprogramming of 3D genome structures in diabetic endothelial dysfunction associated with JUNB.

To further elucidate how JUNB downregulation disrupts chromatin architecture in diabetes, we examined representative endothelial genes involved in vascular homeostasis and function that exhibit altered E-P interactions in BKS db/db endothelial cells. At the *Kcnj2* locus (**Figure 1O**, **Table S4**), which encoding the Kir2.1 potassium channel essential for endothelial membrane potential and functional hyperemia response (both impaired in diabetes^21^). A ∼240 kb E-P loop connecting a distal intergenic enhancer to the *Kcnj2* promoter was completely lost in db/db. This loop exhibited strong JUNB occupancy and H3K27ac enrichment in Ctrl but was nearly absent in diabetes. Similar patterns of disrupted E-P interactions were observed at the *Ccnd2* (**Figure S1F**) and *Igf1* (**Figure S1G**) loci: In control endothelial cells, long-range chromatin loops—spanning approximately 180 kb and 170 kb, respectively—connected distal intergenic enhancers to their target promoters. These loops were enriched for JUNB binding and H3K27ac activity. In db/db endothelial cells, these JUNB-associated loops were lost, accompanied by reduced enhancer activity and diminished transcription of *Ccnd2* and *Igf1*, both of which are key regulators of endothelial cell function and vascular homeostasis^22,23^. At the *Zfp521* locus (a key regulator of hematopoietic stem cell self-renewal in adult hematopoiesis^24^) (**Figure 1P**), a hybrid mode of architectural disruption was observed: one intronic E-P loop (∼120 kb) was markedly weakened, while another distal E-P loop (∼320 kb) was completely lost. This loss of chromatin connectivity was accompanied by reduced enhancer activity and diminished *Zfp521* expression.

To further elucidate the role of JUNB in mouse endothelial cell function *in vitro*, we introduced *Junb* knockdown in wild-type endothelial cells (WT-ECs) using an adenoviral approach (WT-sh*Junb*) (**Figure S1H**). Functional assays were subsequently performed to assess cell migration via wound healing (**Figure S1I and S1J**) and transwell assays (**Figure S1K and S1L**), as well as angiogenic capacity using a tube formation assay (**Figure S1M and S1N**). Compared to control cells (WT-Ctrl), WT-sh*Junb* endothelial cells exhibited significant impairment in cell migration and tube formation, indicative of endothelial dysfunction. These findings highlight JUNB as a key regulator of endothelial homeostasis and suggest that its downregulation may contribute to diabetes-associated vascular impairment. In summary, our results demonstrate that JUNB plays a critical role in regulating enhancer activity and maintaining enhancer-promoter interactions in endothelial cells, and that loss of these JUNB-mediated regulatory mechanisms likely drives endothelial cell dysfunction in the diabetic setting.

### JUNB restoration contributes to chromatin architecture remodelling and vascular homeostasis in hyperglycemic endothelial cells

To investigate whether restoring JUNB expression could reverse chromatin and transcriptional changes associated with hyperglycemia, we cultured HUVECs under normal glucose (NG) or high glucose (HG) conditions, and introduced JUNB overexpression (*JUNB*^OE^) under HG. We then profiled 3D chromatin architecture (H3K27ac HiChIP), enhancer activity (H3K27ac ChIP-seq), chromatin accessibility (ATAC-seq), and gene expression (RNA-seq) (**Figure 2A**).

**Figure 2.**
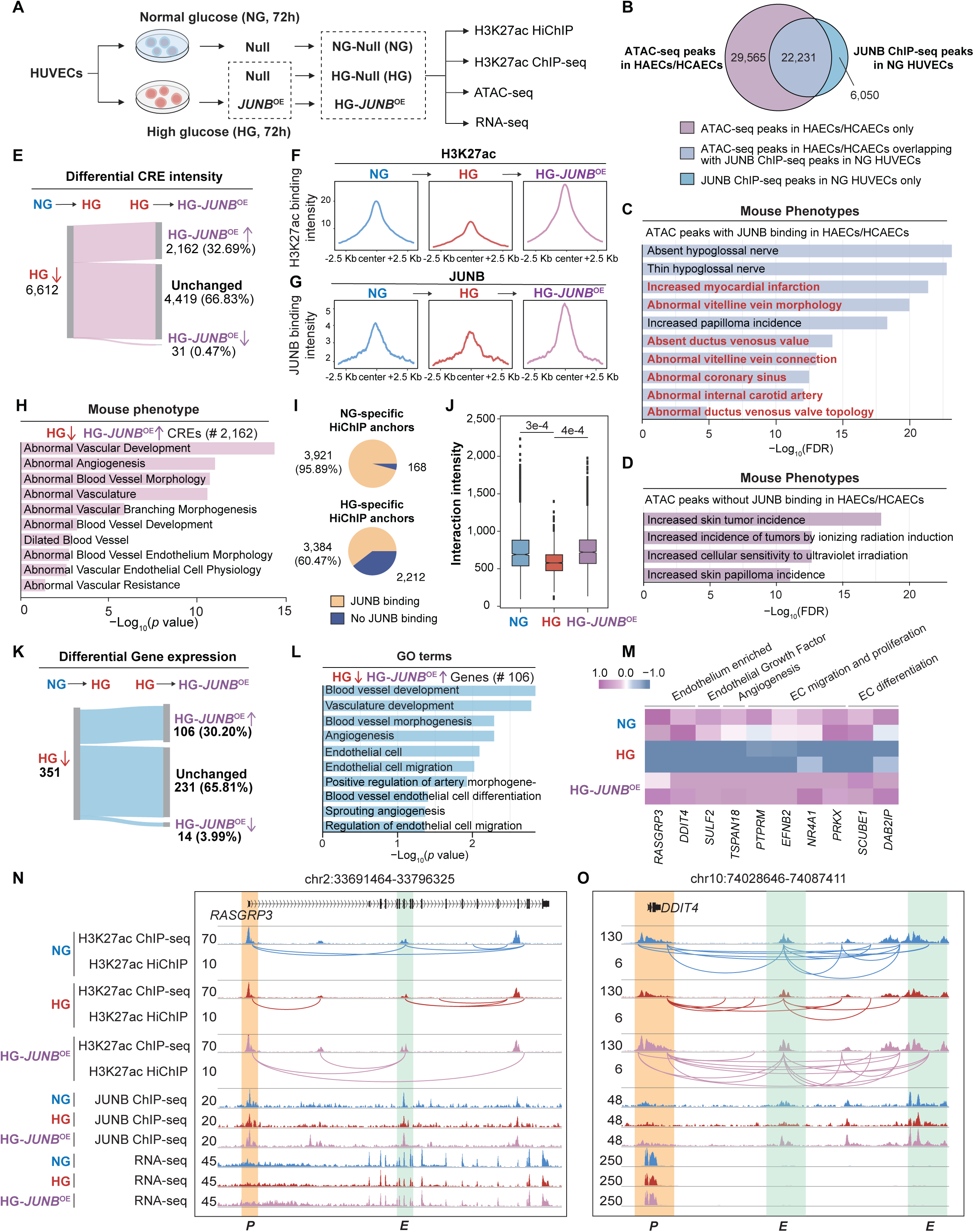
*JUNB* overexpression restores chromatin architecture and angiogenic capacity in hyperglycemic endothelial cells. A. Schematic overview of the experimental design using Human Umbilical Vein Endothelial Cells (HUVECs). HUVECs were cultured in EGM™-2 Endothelial Cell Growth Medium under normal glucose (NG; 5.5 mmol/L D-glucose) or high glucose (HG; 25 mmol/L D-glucose) conditions. An additional HG group was subjected to JUNB overexpression (HG-JUNBOE). Multi-omics datasets, including H3K27ac HiChIP, H3K27ac ChIP-seq, RNA-seq, and ATAC-seq, were generated at 72 hours post-treatment. B. Venn diagram showing the overlap of JUNB binding sites in chromatin accessible regions in primary human endothelial cells. The purple region represents ATAC-seq peaks without JUNB binding in primary human aortic or coronary artery endothelial cells (HAECs/HCAECs); the blue region represents JUNB ChIP-seq peaks in NG HUVECs without ATAC-seq peaks in HAECs/HCAECs. C-D. Bar plots showing the enrichment of “Mouse Phenotype” items enriched in regions overlapping with JUNB binding sites (C) and regions without JUNB binding (D) in HAECs/HCAECs. E. Sankey diagram illustrating the numbers of downregulated CREs under HG compared to NG conditions, and their changes following JUNB overexpression (HG-*JUNB^OE^*). F-G. Aggregation plot showing the H3K27ac (F) and JUNB (G) ChIP-seq signals at downregulated CREs in NG, HG, and HG-*JUNB*^OE^. The signals in a ±2.5 kb window flanking the peak centre were shown. H. Mouse phenotype enrichment analysis of downregulated CREs under HG that were restored by JUNB overexpression (HG-*JUNB*^OE^). I. Proportion of loop anchors specific to NG or HG that were JUNB-bound versus non-JUNB-bound. J. Changes in interaction strength of CREs that were first downregulated from NG to HG and then restored in HG-*JUNB*^OE^. Data were analyzed by Wilcoxon rank sum test with continuity correction. K. Sankey diagram illustrating the numbers of downregulated genes under HG compared to NG conditions, and their changes following JUNB overexpression (HG-*JUNB^OE^*). L. GO enrichment analysis of the genes initially downregulated under HG compared to NG treatment but restored by JUNB overexpression (HG-*JUNB*^OE^). M. Examples of genes that were initially downregulated under HG compared to NG treatment and then restored in HG-*JUNB^OE^*. N-O. Genome browser view showing H3K27ac ChIP-seq, H3K27ac HiChIP, JUNB ChIP-seq and RNA-seq signals at the *RASGRP3* (N) and *DDIT4* (O) loci in NG and HG. Promoters (orange) and enhancers (green) were highlighted.

We first examined whether JUNB binding sites overlapped with chromatin-accessible regions in primary human endothelial cells. Cross-referencing ATAC-seq peaks from HUVECs with those from human coronary artery endothelial cells (HCAEC) and human aortic endothelial cells (HAEC) revealed that a substantial portion (22,231 peaks) of JUNB-bound sites were conserved across primary endothelial cell types (**Figure 2B**, **Figure S2A**), suggesting a shared regulatory landscape and functional relevance of JUNB across vascular beds. To assess the functional relevance of these regulatory elements, we performed GREAT analysis^25^ (mouse phenotype) using the ATAC-seq peaks with or without JUNB binding. The ATAC-seq peaks with JUNB binding were significantly enriched for vascular and cardiovascular phenotypes such as increased risk of myocardial infarction and coronary artery malformation (**Figure 2C**). In contrast, ATAC peaks without JUNB binding were enriched for non-endothelial phenotypes, including increased skin tumor incidence and radiation-induced tumor formation (**Figure 2D**). These results indicate that JUNB preferentially engages regulatory regions functionally associated with endothelial and vascular biology, underscoring its role in maintaining human vascular homeostasis.

We next asked whether *JUNB*^OE^ could rescue chromatin and transcriptional changes induced by HG. Differential analysis of H3K27ac ChIP-seq data revealed that 6,612 CREs were downregulated in HG relative to NG. Remarkably, *JUNB*^OE^ restored H3K27ac signal at 2,162 of these regions (32.7%), while only 31 peaks (0.5%) were further repressed (**Figure 2E**). Aggregate signal profiles centered on CREs further supported this finding: both H3K27ac and JUNB binding intensities were substantially reduced under HG but were reinstated to near-NG levels upon *JUNB*^OE^ (**Figure 2F and 2G**). To assess the functional relevance of these reactivated CREs, we performed GREAT analysis (mouse phenotype) using the 2,162 restored enhancers. Strikingly, these loci were significantly associated with vascular-specific phenotypes, including abnormal vascular development, angiogenesis, blood vessel morphology, and endothelial physiology (**Figure 2H**). These findings demonstrate that JUNB re-engagement at enhancers is sufficient to restore chromatin activity and regulatory programs critical for maintaining vascular integrity under HG stress.

To evaluate whether JUNB overexpression also reinstates 3D chromatin architecture impaired by HG, we analyzed enhancer–promoter interactions via H3K27ac HiChIP. Among NG-specific chromatin loops, 95.9% were marked by JUNB binding, while only 60.5% of HG-specific loops showed JUNB occupancy (**Figure 2I**), indicating a substantial loss of JUNB-associated interactions under HG. Notably, interaction intensity was significantly reduced in HG but was largely rescued upon *JUNB* overexpression, approaching levels observed in NG (**Figure 2J**). These results demonstrate that JUNB is critical for maintaining 3D chromatin structure and that its restoration can reverse HG-induced loss of enhancer–promoter connectivity.

At the transcriptional level, *JUNB* overexpression upregulated 106 of 351 HG-repressed genes (30.20%) while only 14 (3.99%) were further downregulated, indicating broad gene recovery (**Figure 2K**, **Table S5**). These genes were enriched for vascular processes, including endothelial proliferation, migration, differentiation, and sprouting angiogenesis (**Figure 2L and 2M**). Conversely, *JUNB* knockdown under NG suppressed 228 genes linked to similar pathways (**Figure S2B**), underscoring the essential role of JUNB in maintaining endothelial transcriptional programs.

At the *RASGRP3* locus, HG reduced H3K27ac enrichment, JUNB occupancy on the enhancer, and enhancer–promoter connectivity, leading to transcriptional repression. Remarkably, JUNB overexpression restored all three features, reinstating *RASGRP3* expression (**Figure 2N**). At the *DDIT4* locus, HG induced a hybrid pattern of 3D chromatin disruption, characterized by the complete loss of an enhancer–promoter loop and reduced intensity of another enhancer-promoter loop, along with diminished JUNB binding and H3K27ac levels. These changes coincided with transcriptional repression of *DDIT4*. JUNB overexpression effectively restored both chromatin looping and enhancer activity, leading to reactivation of *DDIT4* expression (**Figure 2O**). These two genes were further validated by single-cell transcriptomic data^26^, which showed their preferential expression in vascular endothelial cell clusters across multiple human tissues (**Figure S2C and S2D**).

Consistent with the mouse study, we found that HG exposure impaired endothelial function, while *JUNB* overexpression restored it. Under HG conditions, endothelial cells showed reduced JUNB expression, impaired migration (wound healing and transwell assays), and diminished angiogenic capacity (tube formation assay). Notably, JUNB overexpression rescued these defects, restoring migratory ability and network formation to levels comparable to NG controls (**Figures S2E–S2L**). These results support the conclusion that JUNB is both necessary and sufficient for maintaining endothelial function under diabetic stress via restoring both chromatin architectures and transcriptional activity.

### RBBP6 mediates JUNB degradation and reduces its binding to CREs of key endothelial genes

To elucidate the mechanism underlying JUNB downregulation in diabetic endothelial cells, we first evaluated whether its repression occurs at the transcriptional level. JUNB mRNA levels were not significantly altered in aortic endothelial cells isolated from BKS db/db mice (**Figure S3A**) or in HUVECs treated with HG compared to NG (**Figure S3B**), suggesting that the reduction in JUNB protein under diabetic conditions likely results from post-transcriptional regulation. To identify candidate regulators of JUNB protein stability, we employed rapid immunoprecipitation mass spectrometry of endogenous proteins (RIME)^27–31^ in HUVECs cultured under NG and HG conditions (**Figure 3A**). Quantitative analysis from two independent RIME replicates identified 125 proteins under NG and 116 under HG treatment, including JUNB (**Figure S3C and S3D**, **Table S6**). The relative expression ratio of each enriched protein under NG or HG treatment was determined by dividing its spectral count by that of JUNB under the corresponding treatment. The relative abundance of each co-immunoprecipitated protein, normalized to JUNB, was compared between conditions. A total of 68 proteins exhibited a ratio in HG that was at least 0.5 higher than in NG (**Figure S3E**). Among these, RBBP6, an E3 ubiquitin ligase, was strongly enriched under HG and known to mediate proteasomal degradation of target proteins, implicating it as a potential mediator of JUNB downregulation in diabetes (**Figure 3B**). Immunohistochemical analysis of a human cohort revealed significantly elevated RBBP6 expression in glomerular endothelial cells from diabetic patients (DM, n=84) compared to non-diabetic individuals (NDM, n=90) (**Figure 3C and 3D**). This pattern contrasted with our earlier observation of reduced JUNB protein levels in human ECs (**Figure 1H and 1I**). The differential expression of JUNB and RBBP6 in primary aortic ECs of db/db mice further supported the potential involvement of RBBP6 in JUNB suppression in diabetes(Figure 3E and 3F).

**Figure 3.**
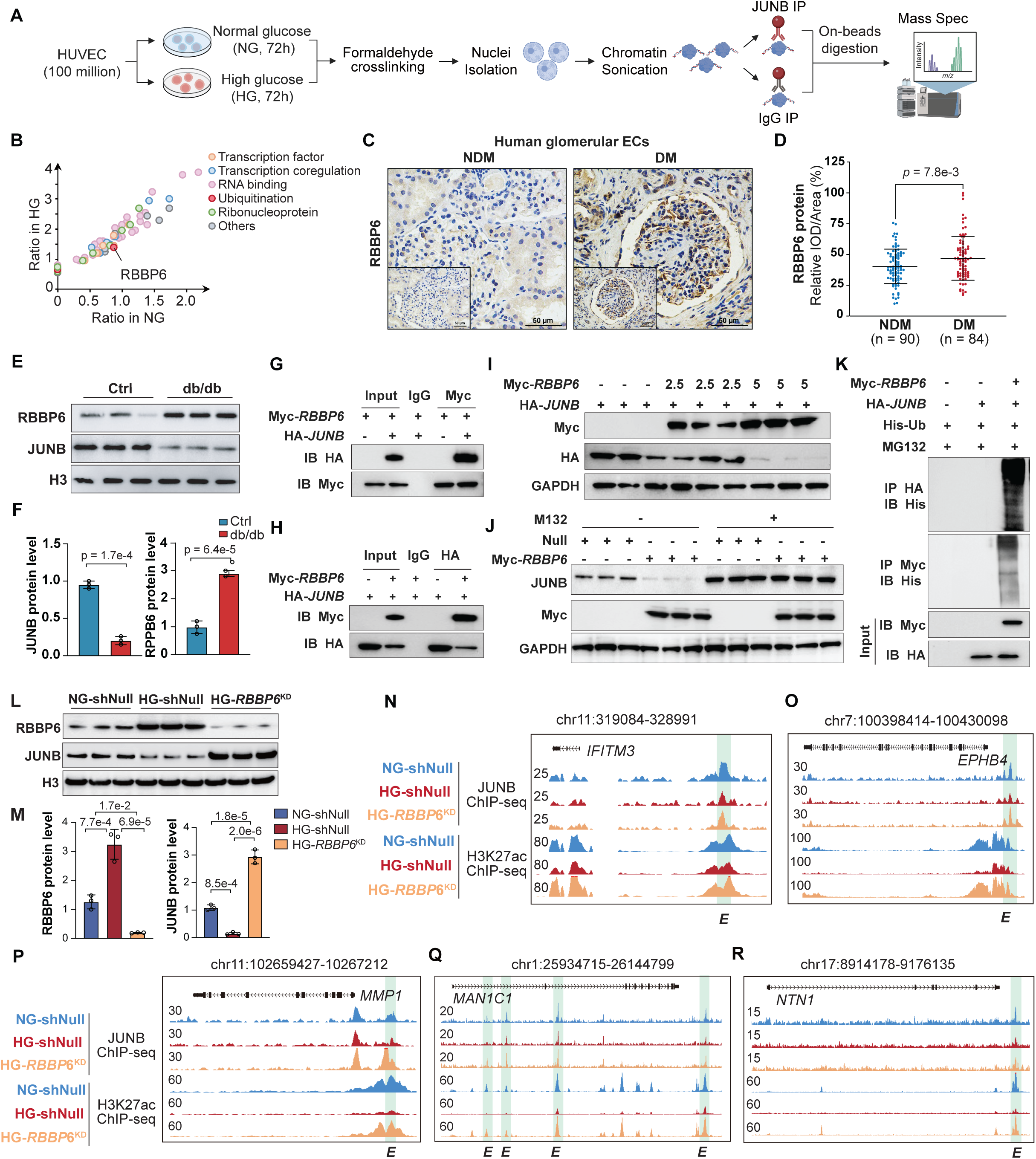
RBBP6 mediates JUNB degradation and reduces its binding to CREs of key endothelial genes. A. Schematic overview of the Rapid Immunoprecipitation Mass Spectrometry of Endogenous Proteins (RIME) procedure. HUVEC cells were cultured under NG or HG conditions for 72 hours before cross-linking, nuclei extraction, lysate sonication, immunoprecipitation and LC-MS/MS analysis. B. Scatter plot showing the relative expression ratio of each enriched protein under HG vs NG treatment. The ratio of each protein was determined by dividing its spectral count by that of JUNB under the corresponding condition. C. Representative immunohistochemical staining of RBBP6 in human glomerular endothelial cells from NDM and DM patients. D. Quantification data analysis of RBBP6 in human glomerular endothelial cells from DM (n = 84) and NDM (n = 90) patients. Data were analyzed by Student’s *t* test. E. Western blot analysis of RBBP6 and JUNB in primary aortic endothelial cells of Ctrl and db/db mouse. F. Quantitative analysis of RBBP6 and JUNB by western blotting in Ctrl (n = 3) and db/db (n = 3) mouse primary aortic endothelial cells. Data were analyzed by Student’s t test. G-H. Co-immunoprecipitation (Co-IP) assay demonstrating the interaction between Myc-RBBP6 and HA-*JUNB* was analyzed by. Following co-transduction with Myc-RBBP6 overexpression and HA-*JUNB* adenovirus, the proteins extracted from HUVECs were immunoprecipitated with anti-Myc antibody (upper panels) or anti-HA antibody (lower panels), respectively (n = 3). I. Western blot analysis of the interaction between Myc-*RBBP6* and HA-*JUNB* in HUVECs transfected with the HA-JUNB adenovirus together with varying amounts of Myc-RBBP6 or empty control adenovirus (n = 3). J. Western blot analysis illustrating the effect of ubiquitin-proteasome inhibitor MG132 on rescuing JUNB protein levels in the presence of Myc-*RBBP6*. K. HUVECs were co-transduced with Myc-*RBBP6*, HA-*JUNB*, and His-*Ub* (ubiquitin), followed by immunoprecipitation with anti-HA (upper panel) or anti-Myc antibodies (middle panel) (n = 3). L. Western blot analysis of RBBP6 and JUNB in primary HUVECs treated with normal glucose (NG-shNull) or high glucose (HG-shNull) and transduced with a null adenovirus, as well as in high glucose with *RBBP6* knockdown (HG-*RBBP6*^KD^). M. Quantitative analysis of RBBP6 and JUNB protein level by western blotting in NG-shNull (n = 3), HG-shNull (n = 3) and HG-*RBBP6*^KD^ (n = 3) treated HUVECs. Data were analyzed by One-way ANOVA (n = 3). N-R. Genome browser views of *MMP1, EPHB4, IFITM3, MAN1C1* and *NTN1* loci. The ChIP-seq peaks of JUNB and H3K27ac in NG-shNull, HG-shNull and HG-*RBBP6*^KD^ HUVECs were shown for each gene. Enhancers were highlighted (green).

To test whether RBBP6 physically interacts with JUNB, we performed reciprocal co-immunoprecipitation assays in HUVECs co-expressing Myc-tagged RBBP6 and HA-tagged JUNB. JUNB robustly co-immunoprecipitated with RBBP6 in both directions (**Figures 3G and 3H**), confirming a direct physical interaction. Then, HUVECs were co-transduced with adenovirus expressing Myc-RBBP6 and HA-JUNB to ectopically co-expressing Myc-RBBP6 and HA-JUNB, and we found that increasing doses of RBBP6 led to a dose-dependent reduction in JUNB protein levels (**Figures 3I**). To explore whether RBBP6 negatively regulates JUNB through ubiquitin-proteasome degradation pathway, HUVECs expressing Myc-RBBP6 were treated with the proteasome inhibitor MG132. Western blot analysis revealed that MG132 treatment significantly attenuated the inhibitory effect of Myc-RBBP6 on JUNB protein expression (**Figure 3J**). To further investigate whether RBBP6 promotes JUNB degradation through ubiquitination, HUVECs were co-transduced with Myc-RBBP6, HA-JUNB, and His-ubiquitin (His-Ub). The cells expressing Myc-RBBP6 exhibited a significant increase in HA-JUNB ubiquitination compared with the cells transduced with a null adenovirus (**Figure 3K**), confirming that RBBP6 promotes JUNB degradation through ubiquitination.

Next, we investigated whether *RBBP6* knockdown could restore JUNB occupancy at CREs of endothelial genes repressed under HG conditions. HUVECs were transduced with adenovirus (NG-shNull, HG-shNull, HG-*RBBP6*^KD^). Western blot analysis showed that JUNB expression increased in RBBP6-knockdown HUVECs, accompanied by decreased RBBP6 protein level (Figure 3L and 3M).

According to the JUNB/H3K27ac ChIP-seq analysis, we observed markedly reduced JUNB binding at enhancer regions of key endothelial regulatory genes under HG condition, including interferon induced matrix metalloproteinase-1 (*MMP-1*), which promotes *VEGFR2* expression and endothelial cell proliferation^32^ (**Figure 3N**); EPH receptor B4 (*EPHB4*), essential for sprouting angiogenesis, vascular morphogenesis, and arteriovenous differentiation^33^ (**Figure 3O**); transmembrane protein3 (*IFITM3*), a pro-angiogenic factor in glioblastoma^34^ (**Figure 3P**). Similarly, the binding of JUNB at enhancers of mannosidase alpha class 1c member 1 (*MAN1C1*), a critical factor for endothelial cell adhesion^35^ (**Figure 3Q**), and netrin-1 (*NTN1*), a key endothelial-derived regulator of angiogenesis^36^ (**Figure 3R**), was also diminished under HG. Notably, knockdown of RBBP6 in HG-treated cells partially rescued both JUNB binding and H3K27ac enhancer activity at these loci, restoring them toward levels observed under NG conditions. These findings demonstrate that RBBP6 contributes to JUNB degradation and its epigenomic disengagement in diabetic endothelial cells, thereby impairing the regulatory networks that sustain vascular homeostasis.

### RBBP6 knockdown restores enhancer-promoter connectivity and transcription of vascular genes to improve endothelial function

To further elucidate the functional consequences of RBBP6 depletion on JUNB-mediated chromatin and transcription regulation, we performed H3K27ac HiChIP and RNA-seq in HUVECs under the three conditions (NG-shNull, HG-shNull, HG-*RBBP6*^KD^), and integrated these data with corresponding JUNB and H3K27ac ChIP-seq profiles. Gene ontology analysis revealed that genes rescued by *RBBP6* knockdown in HG-treated HUVECs were predominantly involved in angiogenesis and vascular permeability (**Figure 4A**). Several key endothelial genes—including *MMP1, EPHB4*, *IFITM3, MANSC1 and NTN1* (**Figure 4B**, **Table S7**)—were transcriptionally repressed in HG-shNull cells but reactivated in HG-*RBBP6*^KD^ cells, accompanied by restored H3K27ac enrichment and JUNB binding at their regulatory elements (**Figure 3N-3R**). Notably, *MANSC1* and *ZDHHC6* also exhibited this rescue pattern and were further validated by single-cell transcriptomic data^26^, which showed their preferential expression in vascular endothelial cell clusters across multiple human tissues (**Figure S4A and S4B**), highlighting their physiological relevance. Locus-specific analysis revealed that *RBBP6* knockdown reinstated the CRE activity at the *MANSC1* locus, with corresponding recovery of JUNB occupancy and enhancer–promoter chromatin looping (∼200 kb) (**Figure 4C**). A similar regulatory architecture was observed at the *ZDHHC6* locus, where *RBBP6* depletion restored JUNB binding, chromatin interactions (∼65 kb), and gene expression (**Figure 4D**).

**Figure 4.**
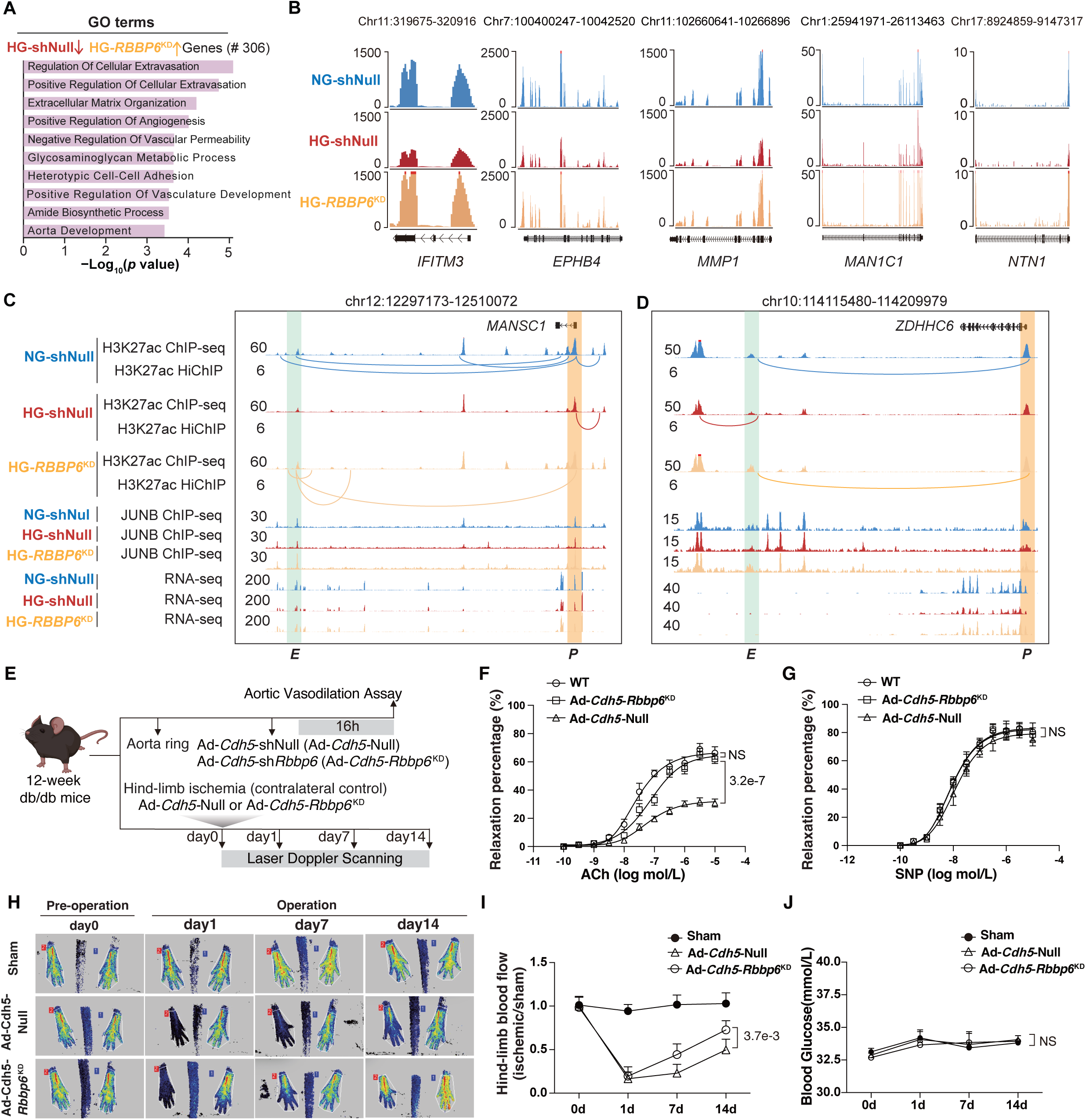
*RBBP6* knockdown restores enhancer-promoter connectivity and transcription of vascular genes to improve endothelial function. A. GO enrichment analysis of genes downregulated in HG versus NG conditions that were restored upon RBBP6 knockdown (HG-*RBBP6*^KD^) in HUVECs. B. Genome browser view showing gene expression profile at the *MMP1, EPHB4, IFITM3, MAN1C1* and *NTN1* loci in NG-shNull, HG-shNull and HG-*RBBP6*^KD^ treated HUVECs. C-D. Genome browser views presenting multi-omics datasets including H3K27ac ChIP-seq, H3K27ac HiChIP, JUNB ChIP-seq, and RNA-seq signals at the *MANSC1* (C) and *ZDHHC6* (D) loci in NG-shNull, HG-shNull and HG-*RBBP6*^KD^ conditions. E. Schematic diagram of mouse hind-limb ischemia model and isometric force measurement procedures. F. Concentration-response curves for acetylcholine (Ach)-induced endothelium-dependent aortic vasodilation in aortas isolated from WT mice or db/db mice (n = 6) treated with Ad-*Cdh5*-Null and Ad-*Cdh5*-*Rbbp6*^KD^. Data were analyzed by Student’s *t* test. G. Concentration-response curves for sodium nitroprusside (SNP)-induced endothelium-independent aortic vasodilation in aortas isolated from WT mice or db/db mice (n = 6) treated with Ad-Null and Ad-*Rbbp6*^KD^. Data were analyzed by Student’s *t* test. H. Representative laser Doppler scanning images of hindlimb ischemia models in db/db mice following different treatments. Ad-*Cdh5*-Null or Ad-*Cdh5*-*Rbbp6*^KD^ was intramuscularly administrated into the ischemic limb immediately after femoral artery ligation (FAL) in db/db mice (n = 7). I. Quantification of hindlimb blood perfusion in db/db mice. Data were analyzed by Two-way ANOVA. J. Quantification of blood glucose levels in db/db mice. Data were analyzed by Two-way ANOVA.

To determine whether *RBBP6* knockdown could mitigate diabetic endothelial dysfunction, we performed *in vitro*, *ex vivo* and *in vivo* functional assays. *In vitro*, *RBBP6* knockdown resulted in enhanced cell migration compared with HG-shNull (**Figure S4C-S4F**). And HUVECs exhibited significantly enhanced tube formation compared to HG-shNull cells after *RBBP6* knockdown, approaching levels observed under NG conditions (**Figure S4G and S4H**). To validate the functional impact of *Rbbp6* knockdown on aortic endothelial function without affecting other cell types, we generated *Rbbp6* shRNA adenovirus vectors with endothelial-specific *Cdh5* promoter (Ad-*Cdh5*-shRNA-*Rbbp6*, hereafter Ad-*Cdh5-Rbbp6*^KD^) and applied it to aortas isolated from db/db mice. Aortic rings were transduced with either Ad-*Cdh5-Rbbp6*^KD^ or Ad-*Cdh5*-shRNA-Null (Ad-*Cdh5-*Null), followed by assessment of vascular reactivity (**Figure 4E**). Endothelium-dependent relaxation in response to acetylcholine (ACh) was significantly improved in Ad-*Cdh5-Rbbp6*^KD^-treated vessels compared to Ad-*Cdh5-*Null (**Figure 4F**), whereas endothelium-independent relaxation to sodium nitroprusside (SNP) remained unchanged between groups **(Figure 4G**). These results indicate that endothelial-specific *Rbbp6* knockdown effectively restores endothelial function in diabetic mouse arteries. To functionally validate the role of endothelial Rbbp6 in diabetic vasculopathy in vivo, a hindlimb ischemia model (femoral artery ligation, FAL) was employed. Immediately after FAL, db/db mice received intramuscular injections of Ad-*Cdh5-Rbbp6*^KD^ or Ad-*Cdh5-*Null into the ischemic limb. Laser Doppler imaging demonstrated markedly improved blood flow recovery in Ad-*Cdh5-Rbbp6*^KD^-treated limbs at days 7 and 14 post-surgery, compared to Ad-*Cdh5-*Null-treated mice (**Figure 4H and 4I**). Importantly, systemic blood glucose levels remained unchanged across groups (**Figure 4J**), indicating that the vascular benefits of *Rbbp6* knockdown are independent of glycemic control. These findings establish endothelial Rbbp6 as a critical regulator of vascular repair and function in diabetic ischemic conditions.

Our results demonstrate that *RBBP6* knockdown not only reinstates JUNB binding and chromatin interactions at vascular genes but also functionally rescues endothelial cell behavior and vascular repair under diabetic stress. These findings establish the RBBP6–JUNB axis as a critical regulator of endothelial homeostasis and a potential therapeutic target in diabetic vasculopathy.

## Discussion

Although transcriptional dysregulation is increasingly recognized as a key factor in vascular complications of diabetes, the chromatin-level transcriptional regulation mechanisms linking metabolic stress to endothelial dysfunction remain poorly understood. A critical unanswered question in the field has been how hyperglycemia perturbs 3D genome organization in endothelial cells and whether specific transcription factors orchestrate these chromatin changes.

In this study, we presented the first high-resolution chromatin architecture atlas in diabetic endothelial cells, generated using H3K27ac HiChIP alongside complementary multi-omic datasets. By profiling several diabetic models, including type 1 and type 2 diabetic mice and human patient-derived endothelial cells, we uncovered widespread rewiring of enhancer–promoter (E–P) interactions induced by hyperglycemia. These architectural changes predominantly occured at JUNB-bound cis-regulatory elements and coincide with transcriptional silencing of key endothelial genes such as *Wnt5a*, *Igf1*, *Zfp521*, and *Kcnj2*. Our data revealed that disruption of JUNB-anchored E–P interactions is a central feature of diabetic endothelial dysfunction.

Previous work has shown that *Junb* is essential for embryonic and retinal vascular development, and that endothelium-specific deletion of *Junb* disrupts vessel formation and angiogenic signaling *in vivo*^37^. However, its role in maintaining chromatin architecture in mature endothelial cells, particularly under metabolic stress, has not been explored. Our results bridge this gap, demonstrating that JUNB not only promotes endothelial gene expression but also protects 3D genome organization. Under hyperglycemia, JUNB protein—but not its mRNA—is selectively depleted. This depletion leads to two modes of chromatin disruption: (1) complete loss of long-range E–P loops and (2) weakening of pre-existing loops. These changes are accompanied by reduced H3K27ac signal and gene repression, directly linking JUNB occupancy to chromatin loop integrity.

To uncover the upstream mechanism of JUNB depletion under diabetic conditions, we identified RBBP6 as a glucose-responsive E3 ubiquitin ligase that promotes JUNB polyubiquitination and proteasomal degradation. RBBP6 is known to be upregulated in several malignancies, including colon cancer^38^ and glioblastoma^39^, and has been reported to target transcriptional regulators such as ZBTB38^40^ and IκBα^38^ for ubiquitin-mediated degradation. A recent study further implicated RBBP6 in diabetic kidney disease, where it promotes the degradation of ERRα, thereby contributing to mitochondrial dysfunction under hyperglycemia^41^. Our findings extended the role of RBBP6 to the vascular endothelium, demonstrating for the first time that it destabilizes an architectural transcription factor, JUNB, under diabetic stress. This interaction represents a direct mechanistic link between hyperglycemia and large-scale chromatin disorganization. Importantly, *RBBP6* knockdown or *JUNB* overexpression in hyperglycemic endothelial cells restored enhancer acetylation, reestablished enhancer–promoter contacts, and rescued the expression of vasculoprotective genes. Nevertheless, *RBBP6* knockdown did not fully restore all JUNB-dependent transcriptional programs, suggesting additional layers of regulation. Future studies should explore whether JUNB cooperates with other co-factors, epigenetic modifiers, or non-coding RNAs to maintain endothelial identity and function under metabolic stress.

In hyperglycemic endothelial cells, either JUNB overexpression or RBBP6 knockdown rescued key cellular functions, including migration and tube formation. These effects extended beyond in vitro assays: ex vivo, endothelial-specific *Rbbp6* silencing significantly improved endothelium-dependent vasodilation in diabetic aortic rings. In vivo, intramuscular Ad-*Rbbp6*^KD^ administration in a diabetic hindlimb ischemia model significantly enhanced perfusion recovery without affecting systemic glucose levels, highlighting the vascular specificity of this intervention. These results suggested that targeting the RBBP6–JUNB pathway could decouple vascular complications from glycemic control, supporting a precision therapeutic strategy for diabetic vasculopathy. Pharmacological stabilization of JUNB or selective inhibition of RBBP6 may offer promising avenues for restoring endothelial integrity and angiogenic capacity in diabetic settings.

Despite significant advances, several important questions remain. First, the upstream signaling pathways and molecular mechanisms that activate RBBP6 in response to hyperglycemia are currently unknown; elucidating glucose-sensitive regulators and post-translational modifications of RBBP6 will be crucial. Second, *RBBP6* knockdown did not fully restore expression of all JUNB target genes, suggesting additional transcription factors, chromatin remodelers or chromatin-associated RNAs collaborate with JUNB to regulate endothelial function. Employing single-cell, multi-omic approaches across diverse vascular beds could provide deeper insights into these cooperative interactions. Third, while short-term benefits of *RBBP6* knockdown are evident, long-term safety and tissue specificity remain unaddressed. Future studies employing endothelial-specific, inducible *Rbbp6* knockout mouse models or small-molecule RBBP6 inhibitors are necessary to rigorously evaluate therapeutic potential. Finally, whether the RBBP6–JUNB regulatory axis contributes broadly to other diabetic microvascular complications, such as nephropathy or retinopathy, warrants further investigation.

In summary, our study uncovers a previously unrecognized mechanism by which hyperglycemia disrupts 3D genome architecture through proteasomal degradation of JUNB. By identifying the RBBP6–JUNB axis as a central regulator of chromatin topology and endothelial function, we establish a mechanistic and actionable link between metabolic stress and epigenomic reprogramming in vascular disease. These insights lay the foundation for novel epigenome-based therapies targeting vascular complications in diabetes.

## Article Information

### Author contributions

Y. F., L.C. W.H., F.L. and L.J., conceived and designed the study. L.J., W.H., X.Y., R.Z. and W.H. conducted all experiments, while L.C., C.Z. and Q.G. performed bioinformatic analyses of ChIP-seq, ATAC-seq, HiChIP, and RNA-seq data. N.T., S.P., S.S., M.W., G.F., A.G.J., G.Q.,and F.L. provided essential tools and contributed intellectual input to experimental design and data analysis. L.J., Y.F., L.C., W.H., S.Z., Y.L. and R.Z. wrote the manuscript with the help of all co-authors.

## Acknowledgments

This work was funded by National Natural Science Foundation of China (Grant no. 82270241), Natural Science Foundation of Guangdong Province (Grant no. 2023B1515020082) (To L.J.); the Science and Technology Project of Shenzhen (nos.JCYJ20240813104118025) (To X.Y.); the Young Scientists Fund of the National Natural Science Foundation of China (Grant No. 82400340) (To L.C.); Shenzhen Science and Technology Program, Shenzhen, China (Grant No. GJHZ20240218111401002), National Science and Technology Major Project (Grant no. 2023ZD0505902) (To Jin-Song Bian); Natural Science Foundation of China Excellent Young Scientists Fund (Overseas) (Grant No. K241141101), Guangdong Basic and Applied Basic Research Foundation for Distinguished Young Scholars (Grant No. 2024B1515020047), Shenzhen Basic Research General Projects of Shenzhen Science and Technology Innovation Commission (Grant No. JCYJ20230807093514029), National Natural Science Foundation of China (Grant No. 82470452), Natural Science Foundation of Xinjiang Uygur Autonomous Region (Grant No. 2024D01D15), Shenzhen Medical Research Funds (Grant No. B2502036) (To Y.F.); and Center for Computational Science and Engineering at Southern University of Science and Technology. The funder had no role in the design, implementation, analysis, interpretation of the data, approval of the manuscript, and decision to submit the manuscript for publication.

---

**Supplementary Figure 1.**
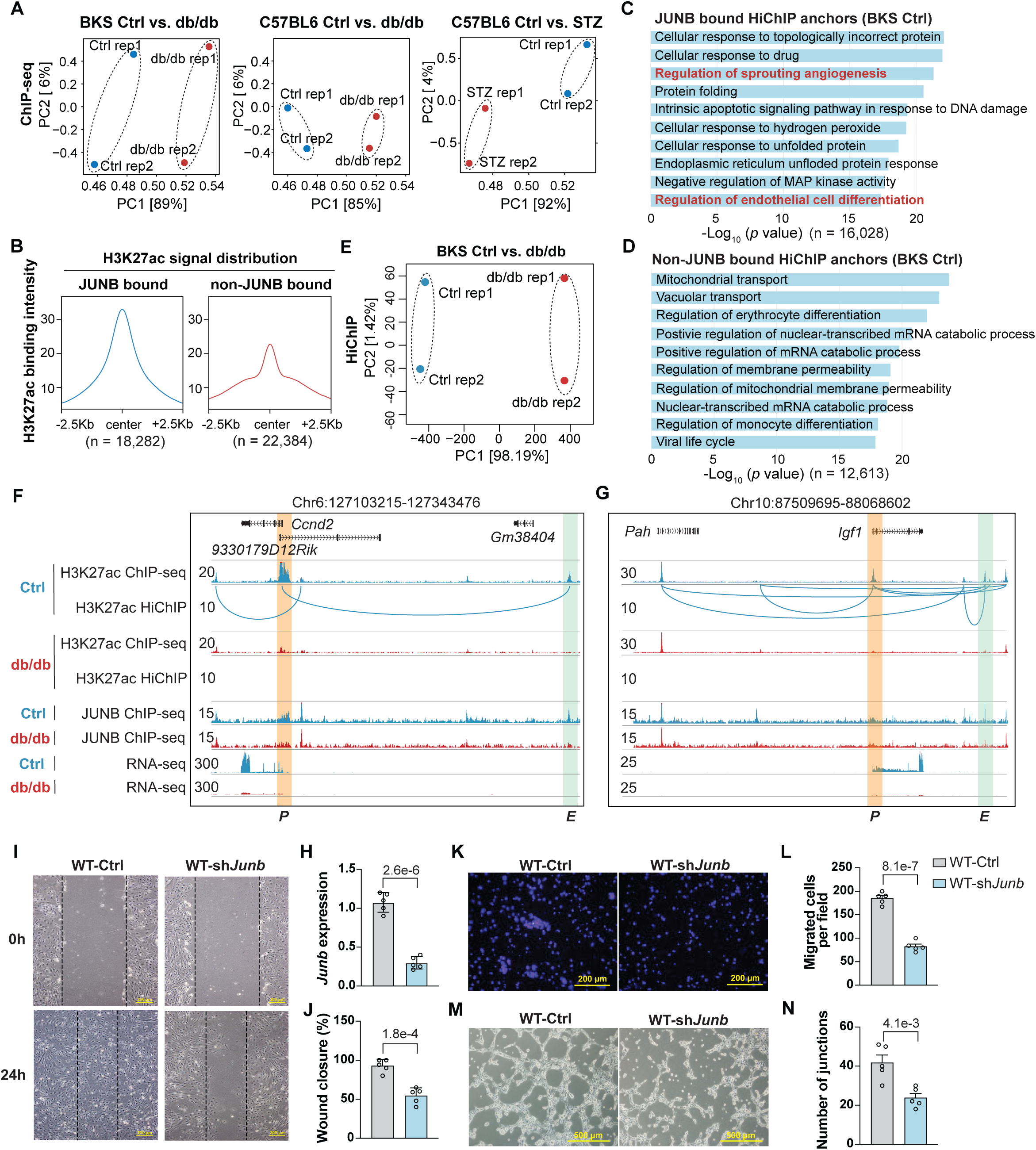

**Supplementary Figure 2.**
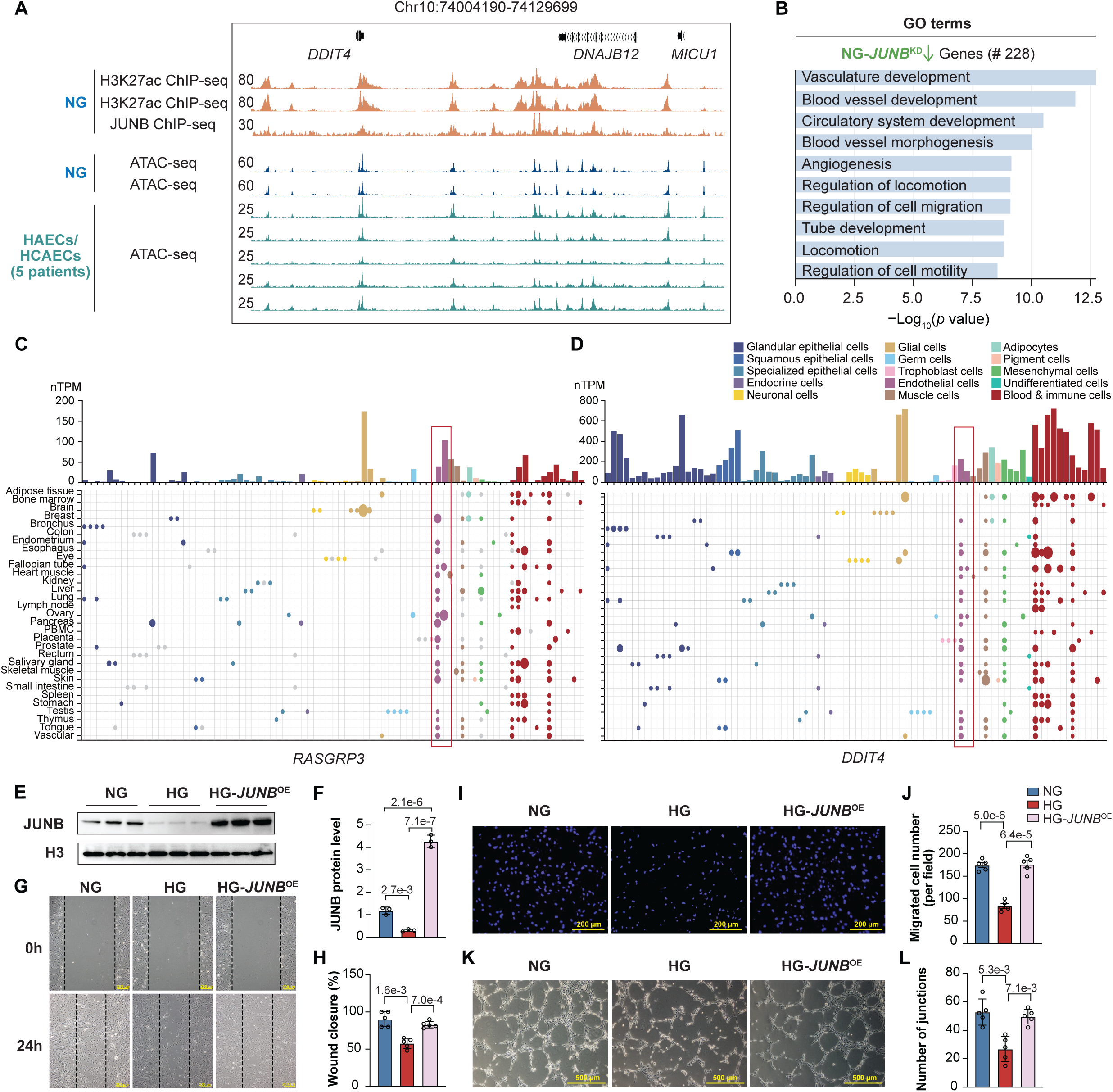

**Supplementary Figure 3.**
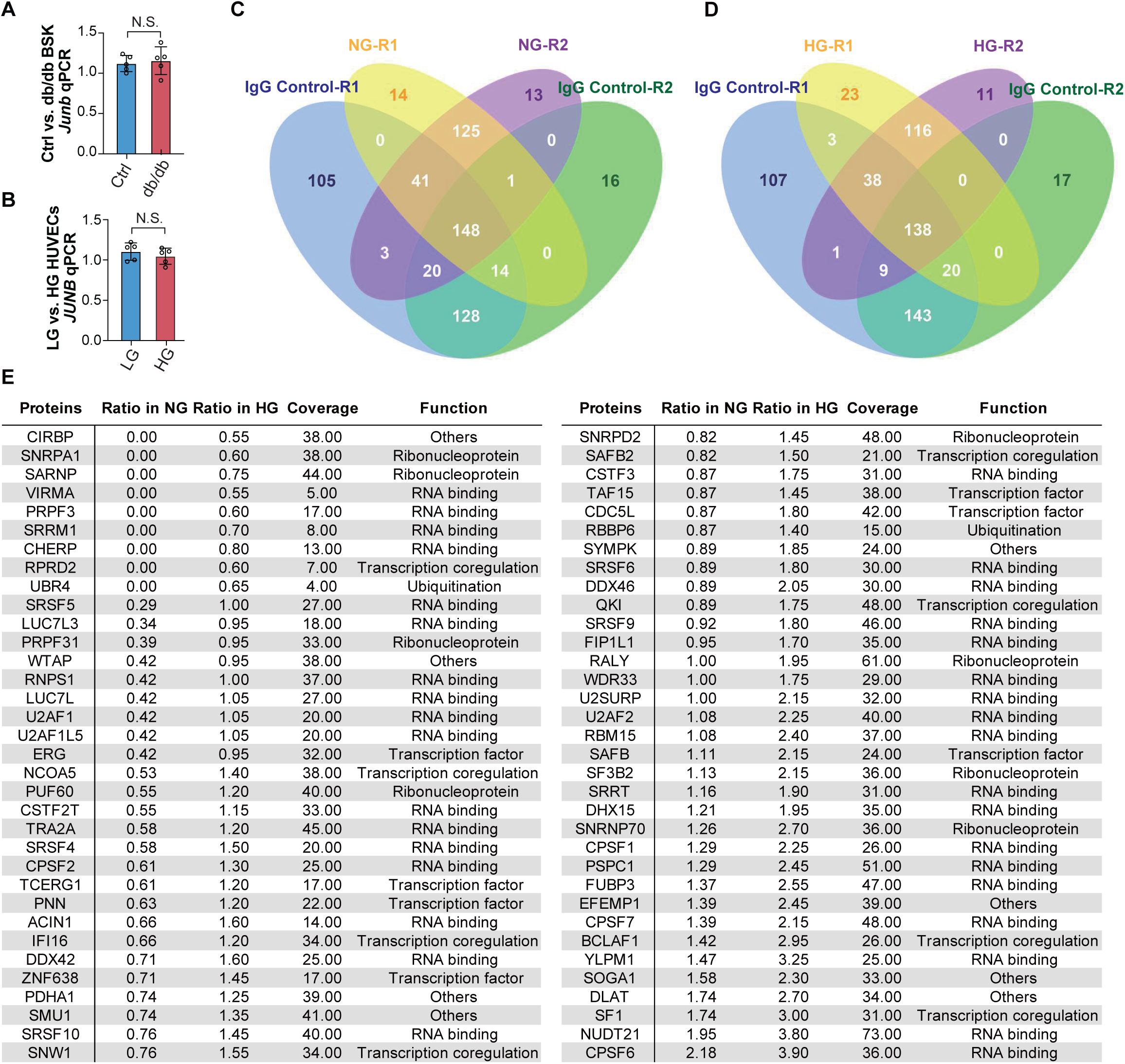

**Supplementary Figure 4.**
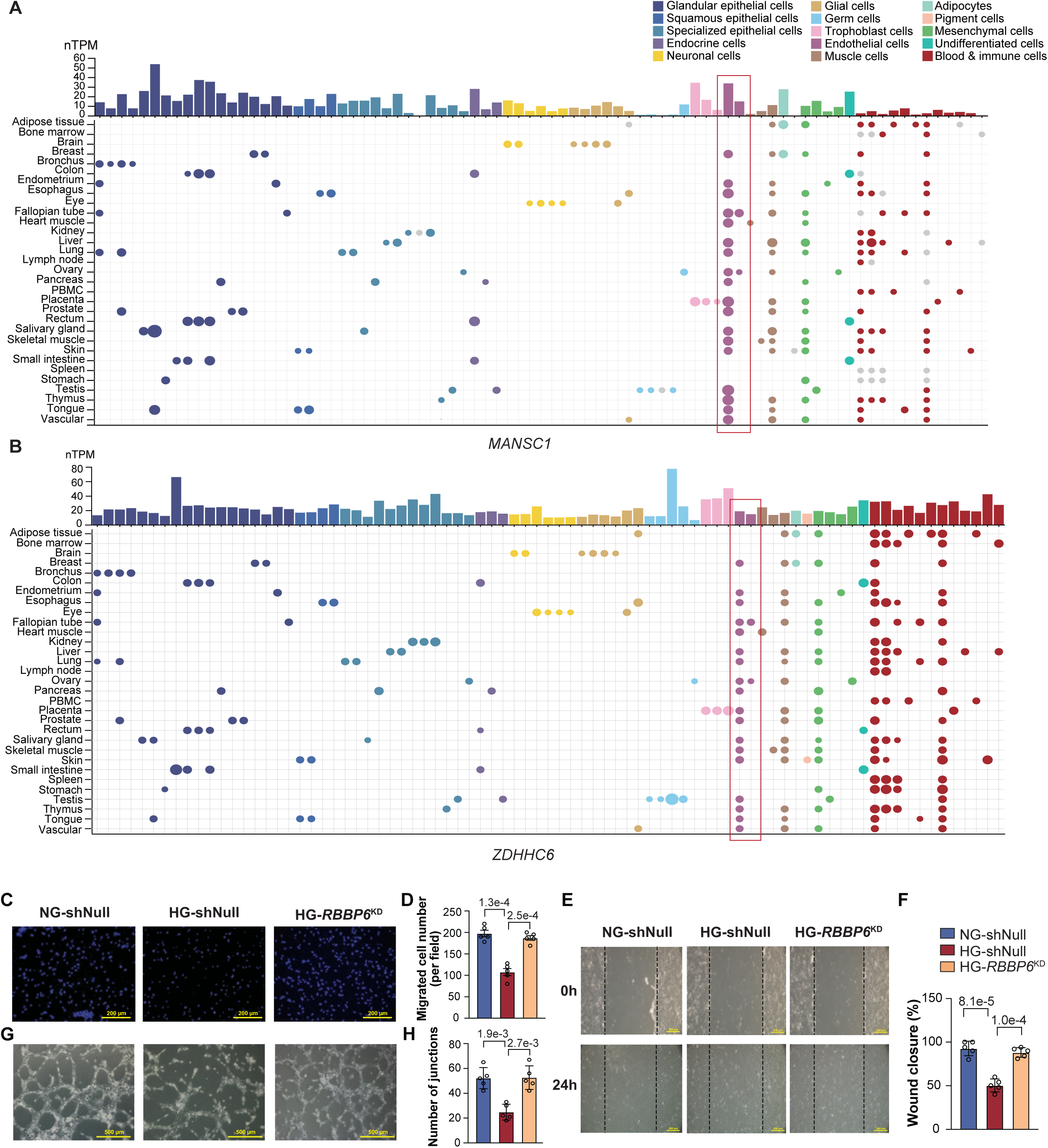

