## Supplementary material for "Rewiring of 3D Enhancer-Promoter Interactome Underlies Diabetic Endothelial Dysfunction": methods and supplementary figure legends

### Methods and Materials

#### Animals

All animal experiments were approved by the Institutional Review Board (or Ethics Committee) of Guangdong Provincial People's Hospital (Protocol code KY2020-131-01). Every effort was made to minimize animal distress and to limit the number of animals involved. All type I, type II (C57BL6 db/db and BKS db/db) and control mice (C57BL6 Ctrl and BKS Ctrl) were purchased from the Jackson Laboratory. All experimental animals were 6-week-old male mice. For type I diabetic mice, daily IP injections of 50 mg STZ/kg body weight was performed on wild type mice (C57BL/6J, six-week-old males) for five consecutive days. Only mice with blood glucose exceeding 250 mg/dL were determined as type I diabetic mice (STZ) used for this study. Age matched controls received the buffer (NaCl) injections.

#### Mouse aortic endothelial cell isolation and culture

After euthanization, the abdominal and thoracic cavities were opened to expose the aorta and heart. Then the abdominal aorta was flushed with 5 ml PBS containing 1,000 U/mL of heparin to remove blood. The thoracic aorta was isolated and cut into the 1-mm rings. The rings were cultured on a matrix pre-coated 6-well plate in the endothelial cell growth medium at 37°C under 5% $CO_2$. After 4 days, the aortic segments were gently removed from the matrix without interrupting the growing endothelial cells. The aortic cells were obtained using neutral proteinase (50 U/mL) and incubated with CD31 MicroBeads (Miltenyi Biotech, 130-097-418) according to the manufacturer's protocol. The magnetically labeled CD31+ cells were resuspended in the endothelial cell growth medium and cultured in gelatin-pre-coated T25 flasks at 37°C under 5% $CO_2$. Cells were generally ready for the experiment after 2–3 passages.

**Human endothelial cell culture**

The human endothelial cells, including Type I diabetic patient-derived aortic endothelial cells (HAEC, CC-2919), Type I diabetic patient-derived coronary artery endothelial cells (HCAEC, CC-2921), Type II diabetic patient-derived aortic endothelial cells (CC-2920), Type II diabetic patient-derived coronary artery endothelial cells (CC-2922), normal human aortic endothelial cells (CC-2535), normal human coronary artery endothelial cells (CC-2585), and human umbilical vein endothelial cells (HUVEC) obtained from Lonza. All cells were cultured in a completed medium (EGM™-2 Endothelial Cell Growth Medium, CC-3162, Lonza) at 37˚C and with 5% CO2. For high glucose (HG) treatment, the cells were cultured in 25 mmol/L D-glucose (MilliporeSigma G8644) for 72 hours. The cells cultured in EGMTM-2 mediums (5.5 mmol/L) for 72 hours were used as normal-glucose (NG) treatment groups.

**Adenovirus transduction**

All the adenovirus was developed by Vector Biolabs. The day before transduction, endothelial cells were seeded at a density of $2×10^5$ cells/well in 6-well plates. 4-6 hours prior to infection, the old medium was aspirated and replaced with fresh medium. When performing infection, the completed medium was replaced with 1 ml pre-mixed virus/basic medium (without growth factors and FBS).

After 2 hours, the pre-mixed virus/basic medium was replaced with the complete medium at 37˚C and with 5% CO2. The wild-type mouse aortic endothelial cells were transduced with control shRNA-Null adenovirus (WT-Ctrl) or shRNA-*JUNB* adenovirus (WT-sh*JUNB*). NG-treated HUVECs were transduced with CMV-Null adenovirus (NG), shRNA-Null adenovirus (NG-shNull), or shRNA-JUNB adenovirus (NG-*JUNB*$^{KD}$). HG-treated HUVECs were transduced with null adenovirus (HG), CMV-JUNB overexpression adenovirus (HG-*JUNB*$^{OE}$), shRNA-Null adenovirus (HG-shNull), or shRNA-*RBBP6* (HG-*RBBP6*$^{KD}$).

**Western blot analysis**

Human or mouse cells were lysed with ice-cold cell lysis buffer (Cell Signaling Technology, 9803S) in the presence of protease inhibitor (Roche, 04693132001). After centrifuging at 12,000 rpm for 20 min at 4 °C, the protein supernatant was transferred into a new cold Eppendorf tube and kept on ice. Following determination of the protein concentration by the Bradford assay (Bio-Rad, 5000001), the protein samples (40 µg) were resolved in 10x NuPAGE reducing agent in the presence of 4x LDS sample buffer and denatured at 70°C for 10 min prior to loading on the NuPAGE™ 4-12% Bis-Tris Protein Gels (Thermo Fisher Scientific, WG1402BOX). Following electrophoresis, the proteins were transferred from the gel to nitrocellulose membranes using semi-dry transfer (Power Blotter Station, Thermo Fisher Scientific, PB0010). Equal loading and transfer of proteins was confirmed by the quantitative Ponceau red staining. The membranes were incubated for 60 min with 5% dry milk and Tris-buffered saline was used to block the nonspecific binding sites. The membranes were immunoblotted overnight at 4 °C with antibodies against JUNB (Cell Signaling Technology, C3753, 1:2000 dilution), JUND (Cell Signaling Technology, 5000, 1:2000 dilution), c-JUN (Cell Signaling

Technology, 9165, 1:2000 dilution), FOS (Cell Signaling Technology, 31254, 1:2000 dilution), anti-Myc-Tag (Cell Signaling Technology, 2276, 1:2000 dilution) and anti-HA-Tag antibody (Cell Signaling Technology, 3724, 1:2000 dilution), and anti-His-Tag (Cell Signaling Technology, 12698 dilution), GAPDH (MilliporeSigma, G9545), or H3 (Cell Signaling Technology, 4499, 1:2000 dilution) on a rocking platform. Following washing for three times for 5 min each with Tris-buffered saline, the membranes were incubated for 60 min with an HRP-conjugated secondary antibody, washed three times with Tris-buffered saline, and developed with the SuperSignal™ West Pico PLUS Chemiluminescent Substrate (Thermo Fisher Scientific, 34580).

**Co-IP assay**

Co-IP was performed according to the kit manual (ThermoScientific, 88804). After cells were transduced with adenovirus carrying Myc-RBBP6, HA-JUNB, and His-Ubiquine for 48 hours, the cells were harvested and lysed with ice-cold cell lysis buffer in the co-IP kit. Anti-Myc, anti-HA, or anti-His magnetic beads were added to the protein samples (200 µg) to pull down Myc-RBBP6, HA-JUNB, and His-Ubiquitin, respectively. The normal mouse IgG (Cell Signaling Technology, 6880, 1:2000 dilution) or normal rabbit IgG (Cell Signaling Technology, 2729, 1:2000 dilution) was used as a negative control. The immunoprecipitants were subjected to Western blotting to analyze the protein expression.

**ChIP-seq on mouse aortic endothelial cells and HUVECs**

10 million cells were collected and washed with PBS twice, then suspended in 10 ml PBS. Cells were fixed with 1% formaldehyde (Thermo Fisher Scientific, 28906) with rotation for 10 minutes and

fixation was quenched with 0.125M glycine (final concentration). Next, cells were washed with cold PBS (with proteinase inhibitors cocktails, PIC) and lysed in SDS lysis buffer. The pellets were suspended in 0.33% SDS incubation buffer (0.3% SDS, 1.6% Triton X-100, 100mM NaCl, 50mM Tris pH 8.0, 5mM EDTA pH 8.0, 1 x PIC) and sonicated to achieve enrichment of 300-500 bp DNA fragments on average. Sonicated DNA was quantitated with Qubit spectrophotometer and checked by Agilent 2100 bioanalyzer.

An aliquot of chromatin (8 µg) was precleared with protein A/G agarose beads (Thermo Fisher Scientific). Specific DNA regions were precipitated by 4 µg of antibody against H3K27Ac (Active Motif, 39133) or 10 µg of antibody again JUNB (Cell Signaling Technology, C3753) with overnight rotation at 4 °C. 30 µl protein A/G magnetic beads were added to each ChIP sample, and incubated with rotation at 4 °C for 2 hours. After appropriate washing, beads were suspended in 200 µl elution buffer (1% SDS, 0.1 M $NaHCO_3$) and incubated for 30 minutes with shaking (900 rpm) at 65 °C. The eluted ChIP supernatant (without beads) was transferred to a new EP tube. 38. NaCl (200 mM final conc) and Proteinase K (final conc, 0.2 mg/ml) was added to each tube and incubated in a thermomixer overnight with 1000 rpm shaking at 65°C to reverse the crosslinks. ChIP DNA was purified using the Qiagen MinElute PCR purification kit and measured with the Qubit. Quantitative PCR was performed to check the ChIP efficacy.

ChIP-Seq libraries were generated from the ChIP-DNA using a custom Illumina library type on an automated system (Apollo 342, Wafergen Biosystems/Takara). ChIP-Seq libraries were sequenced on Illumina NextSeq 500 as 75-nt single end reads. Adapter sequences were not trimmed during demultiplexing.

**H3K27ac HiChIP (mECs and HUVECs)**

H3K27ac HiChIP library preparation was performed by Arima Genomics (https://arimagenomics.com/). Proximally-ligated chromatin labeled with biotin was prepared using the Arima-HiC kit (Arima Genomics, A510008). First, crosslinked chromatin was digested with a specialized restriction enzyme cocktail to create DNA fragments with 5'-overhangs. These overhangs were subsequently filled in and labeled with biotinylated nucleotides, enabling downstream capture. The spatially proximal DNA ends were then ligated. The resulting proximally-ligated chromatin was fragmented using the Diagenode Bioruptor Pico instrument, enriched for H3K27ac modifications by overnight binding to an H3K27ac-specific antibody (Abcam, ab4729) and subsequent immunoprecipitation with Protein A magnetic beads (Thermo, 10002D). Following immunoprecipitation, the crosslinks were reversed, and the chromatin was purified using the Arima-HiC kit. For sequencing library preparation, biotin-enriched chromatin fragments were isolated using streptavidin-coated C1 magnetic beads (Thermo, 65001). The samples were tagmented on-bead utilizing Tagment DNA Enzyme and Buffer (Illumina, 20034197). Subsequently, DNA was ready for PCR enrichment. Tagmented DNA was PCR amplified utilizing NPM and the indexing primers from the Nextera XT DNA Library Prep kit (Illumina, FC-131-1024). The PCR products were purified and size selected by 0.5X SPRI beads followed by 0.7X SPRI beads. The resulting libraries were high-quality and Illumina-compatible, ready for downstream sequencing on Illumina Novaseq 6000.

**ATAC-seq (mECs, HUVECs, normal human aortic endothelial cells and Normal human coronary artery endothelial cells)**

Mouse endothelial cells (mECs) and HUVECs were pretreated with 200 U/ml DNase (Worthington) for 30 min at 37 °C to remove free-floating DNA and to digest DNA from dead cells. This treatment effectively minimizes the interference from exogenous or dead cell DNA and reduces background noise. This medium was then washed out, and the cells were resuspended in cold PBS. After the cells were counted, 50,000 cells were resuspended in 1 ml of cold ATAC-seq resuspension buffer (RSB; 10 mM Tris-HCl pH 7.4, 10 mM NaCl, and 3 mM MgCl2 in water). Cells were centrifuged at 500 g for 5 min at 4 °C. Supernatant was carefully aspirated. Cell pellets were then resuspended in 50 μl of ATAC-seq RSB containing 0.1% NP40, 0.1% Tween-20, and 0.01% digitonin by pipetting up and down three times. This cell lysis reaction was incubated on ice for 3 min. After lysis, 1 ml of ATAC-seq RSB containing 0.1% Tween-20 was added, and the tubes were inverted to mix. Nuclei were then centrifuged for 10 min at 500 g at 4 °C. Supernatant was removed and nuclei were resuspended in 50 μl of transposition mix (25 μl 2× TD buffer (10mM Tris-HCl pH 7.4, 10mM NaCl, 3mM MgCl2), 2.5 μl transposase (100 nM final), 16.5 μl PBS, 0.5 μl 1% digitonin, 0.5 μl 10% Tween-20, and 5 μl water) by pipetting up and down at least five times. Transposition reactions were incubated at 37 °C for 30 min in a thermomixer with shaking at 1,000 r.p.m. DNA fragments were cleaned up with Zymo DNA Clean and Concentrator-5 kit. Libraries were prepared as the description (Buenrostro et al., 2015). Sequencing of mEC libraries was performed on the Illumina NovaSeq 6000 platform, and HUVEC libraries were sequenced using the Illumina NextSeq 500 platform.

**RNA-seq (mECs, HUVECs, normal human aortic endothelial cells and Normal human coronary artery endothelial cells)**

Total RNA was extracted using Trizol (Invitrogen, Carlsbad, CA, USA) according to the manufacturer's instructions. After extracting the total RNA from samples, mRNA and non-coding RNAs were enriched by removing rRNA form the total RNA with kit. By using the fragmentation buffer, the mRNA and non-coding RNAs were fragmented into short fragments (about 200~700 bp), then the first-strand cDNA was synthesized by random hexamer-primer using the fragments as templates. Buffer, dNTPs, RNase H and DNA polymerase I were added to synthesize the second strand cDNA. The double strand cDNA was purified with QiaQuick PCR extraction kit and then used for end-polishing. Sequencing adapters were ligated to the fragments, then the second strand was degraded using UNG(Uracil-N-Glyosylase) finally. The fragments were purified by Agarose gel electrophoresis and enriched by PCR amplification.The library products were ready for sequencing analysis via Illunima HiSeq 4000. The libraries of primary normal human aortic and coronary artery endothelial cells were sequenced on the Hiseq X Ten.

**Mouse vessel preparation and isometric force measurement**

After euthanization, the thoracic aorta was removed and placed in an ice-cold oxygenated Krebs-Henseleit solution. The dissected aorta was cut into several 2-mm-long ring segments and cultured in the plate with DMEM supplemented with 40% serum collected from db/db mice[1]. After infection with Ad-*Cdh5*-shRNA-Null (Ad-Null) or Ad-*Cdh5*-shRNA-*RBBP6* (Ad-*RBBP6*$^{KD}$) for 16 hours, the aortic rings were mounted to a Multi Wire Myograph System (Danish Myo Technology, East Jutland, Denmark) to assess isometric tension. Acetylcholine (ACh; $10^{-10}$–$10^{-5}$ M) or sodium nitroprusside (SNP; $10^{-10}$ –$10^{-5}$ M) were administrated to evaluate endothelium-dependent aortic vasodilation and endothelium-independent aortic vasodilation respectively.

#### Mouse hindlimb ischemia model

The hindlimb ischemia model was established using 10–12-week-old db/db mice as previously described[2]. Under anesthesia, the left femoral artery was exposed and occluded using triple knots to induce ischemia. Ad-Null or Ad-*RBBP6*$^{KD}$ at $2\times10^{11}$ viral genomes (vg) was intramuscularly administrated into the ischemic limb immediately following femoral artery ligation. The contralateral hind limb was sham-operated (Sham) served as an internal control. Blood flow was monitored using laser Doppler scanning analyzer (Moor LDI; Moor Instruments) at baseline (pre-operation) and on post-operative days 1, 7 and 14 to access perfusion changes.

#### Cell migration and invasion assays

The transwell system (8-μm-pore size, 24 well plate, FisherScientific, 07-200-174) was used to analyze cell migration and invasion assays. Endothelial cells were detached by trypsinization and seeded in the upper compartment of Transwell plate ($5\times10^4$ cells/insert) pre-coated with 0.1% gelatin, which was then inserted into a well containing the complete medium. Endothelial cells were allowed to migrate across the membrane for 3 hours, after which the inserts were removed, and the cells were fixed with formaldehyde for 10 minutes. After washing with water, a sterile cotton swab was applied to the top of the insert to scrap off the cells which had not migrated through the membrane. The migrated cells on the lower side of the insert membrane were stained with DAPI (10 μg/ml) in 1% Trition X-100 solution for approximately 10-15 minutes. After washing with PBS, the cells on the membrane were photographed and counted at least three random fields.

Scratch assays were also used to analyze cell migration. Cells were starched with 100μl pipette tips and then cultured in medium for 24 hours. After photography, the wound closure was measured by the distances between the two edges of the scratched wound.

**Tube formation assay**

Endothelial cells were seeded at a density of $3\times10^4$ cells/well in 12-well plates containing 300 μl solidified Matrigel (FisherScientific, CB354248) at 37˚C, 5% CO2 in the cell culture incubator. After 6-hour incubation, each well was photographed. The number of the branch junctions or tubules was counted in at least three individual wells.

**Rapid immunoprecipitation mass spectrometry of endogenous protein (RIME)**

HUVECs were cultured under NG or HG conditions for 72 hours before magnetic beads conjugated with specific antibodies were prepared. To preserve protein-DNA interactions, cells underwent dual cross-linking with 2 mM disuccinimidyl glutarate (DSG) for 20 minutes, followed by 1% formaldehyde (FA) for 10 minutes. Nuclei were then isolated, and chromatin was fragmented via sonication, yielding DNA fragments enriched at approximately 500 bp. The sheared 150 μg chromatin was subsequently incubated with antibody against JUNB (Cell Signaling Technologies, 3753BF) -conjugated beads or IgG-conjugated beads for 16h, followed by rigorous washing steps to reduce nonspecific binding. Immunoprecipitated protein complexes were then subjected to on-bead digestion, and the resulting peptides were analyzed by liquid chromatography-tandem mass spectrometry (LC-MS/MS). Briefly, Active Motif conducted RIME assays on dual cross-linking HUVECs samples.

**ChIP-seq data analysis**

The hg19 (human) and mm10 (mouse) genome primary assembly build were obtained from Gencode database[3]. Single-end reads were mapped to the hg19 (human) or mm10 (mouse) reference genome using the bwa[4] mem algorithm with default parameters. The duplicated reads were identified and removed using MarkDuplicates from the Picard Toolkit (https://broadinstitute.github.io/picard/), and reads with a mapping quality score (MAPQ) lower than 20 were filtered out using samtools[5]. Additionally, reads mapping to mitochondrial DNA and ENCODE blacklisted regions were excluded from further analysis to reduce potential bias. The ChIP-seq peaks were called using MACS2[6] with default narrow peak settings (q-value cutoff = 1e-5, model fold = [10,30], band width = 300 bp). The promoter regions were defined as +/- 2.5Kb around the transcription start sites (TSSs). H3K27ac peaks overlapping with these promoter regions were defined as "proximal promoter peaks" (P), whereas peaks located outside these regions were designated as "distal enhancer peaks" (E).

**Differential ChIP-seq peak analysis**

For differential binding analysis between conditions, ChIP-seq datasets were processed using the DiffBind package (https://bioconductor.org/packages/release/bioc/html/DiffBind.html) in R. An universal peak set was constructed by merging all replicate peak calls, samples using the dba.count function with minOverlap=2, ensuring that any peak present in at least two replicates was included. Peak merging produced a single binding matrix that included all samples. For each peak in this universal set, read counts were quantified for each sample in a standardized 500 bp region (±250 bp) around the peak summit. These per-sample counts were then used to perform differential peak analysis.

Differential binding was then evaluated using the edgeR[7]. Peaks with FDR < 0.05 and fold change > 1 were considered significantly different between conditions. For Principle component analysis (PCA), the variance stabilizing transformation was first applied, and the top 5,000 most variable peaks were selected for visualization. To visualize the relationship between binding intensity and differential binding, MA plots were generated using R package ggplot2, where the M-axis represents the log2 fold change between conditions and the A-axis represents the mean normalized read counts (average binding intensity).

**ATAC-seq data analysis**

Raw FASTQ files were processed to remove adapters using Trimmomatic[8]. The filtered reads were mapped to mm10 using bwa[4] mem with default parameters. Duplicated reads were removed using Picard MarkDuplicates (https://broadinstitute.github.io/picard/), and reads with MAPQ scores below 20 were filtered out using samtools[5]. Additionally, reads mapping to mitochondrial DNA and ENCODE blacklisted regions were excluded from further analysis. Peak calling was performed using MACS2[9] in ATAC-seq mode with parameters: --format BAMPE --nomodel --shift -100 --extsize 200 --keep-dup all -g mm --call-summits -q 0.05. Additionally, genomic coverage files (bigWig format) were generated using deepTools[10] bamCoverage with RPKM normalization for visualization in genome browsers.

**Transcription factor motif enrichment analysis**

Motif analysis was conducted using Homer (http://homer.ucsd.edu/homer/, v4.11). Known motif collection HOCOMOCOv11_full_MOUSE_mono_homer_format_0.0001.motif was downloaded from HOCOMOCO databse[11]. H3K27ac ChIP-seq peaks located outside promoter regions (>2kb from TSS) were intersected with ATAC-seq peaks using BEDtools[12] intersect command, requiring a minimum overlap of 1bp. The corresponding DNA sequences were extracted from the mm10 reference genome using BEDtools[12] getfasta. De novo motif discovery was performed using HOMER findMotifs.pl script with parameters: -len 8,10,12 -size given -S 10 -nomotif -p 8, searching for motifs of lengths 8, 10, and 12 bp. Background sequences were automatically generated by HOMER using random GC-content-matched sequences from the genome. For known motif enrichment analysis, the program was run in known motif discovery mode (-known) against the HOCOMOCO database. Motifs were considered significantly enriched when meeting the criteria of q-value < 0.01 and fold enrichment > 2 compared to background.

**RNA-seq data analysis**

Raw sequencing reads were first subjected to quality control. The reads were mapped to hg19 or mm10 using STAR[13] alinger. Gene-level quantification was performed using featureCounts[14] from the Subread package, with specific parameters set to account for paired-end reads (-p), count at exon features (-t exon), and aggregate counts at gene level (-g gene_id). Gene annotations were obtained from GENCODE. For differential expression analysis, the edgeR package was employed using a robust statistical framework. The analysis pipeline included TMM (Trimmed Mean of M-values) normalization to account for library size differences and composition biases, followed by estimation of dispersion using the robust empirical Bayes method. A generalized linear model (GLM) was fitted to the data, and differential expression was assessed using likelihood ratio tests. Genes with FDR < 0.05 and absolute fold change > 1.5 were considered differentially expressed.

**Analysis of the relationship between H3K27ac signal intensity and JUNB binding**

To investigate the relationship between H3K27ac peak intensity and JUNB occupancy in BKS Ctrl mice, H3K27ac peak regions were categorized into JUNB-bound and JUNB-unbound regions based on the overlap with JUNB ChIP-seq peaks. Each H3K27ac peak region was then extended to a fixed length of 5 kb centered on the peak summit. These regions were then binned into 10 bp intervals. For each bin, H3K27ac ChIP-seq read density was calculated after normalizing using RPKM. An aggregation plot was generated to visualize the average H3K27ac signal distribution across these regions, with the x-axis representing the relative distance from the peak center (-2.5 kb to +2.5 kb) and the y-axis showing the mean normalized H3K27ac signal intensity.

**Analysis of the relationship between down-regulated (BKS Ctrl versus db/db) H3K27ac peaks and JUNB binding**

We classified the down-regulated H3K27ac peaks into three categories based on JUNB binding patterns: regions with reduced JUNB binding, regions with unchanged JUNB binding, and regions without JUNB binding. For each category, we analyzed the signal distribution of three genomic features: JUNB occupancy, H3K27ac occupancy, and chromatin accessibility (ATAC-seq). We extended the genomic region to ±2.5kb around each peak summit and binned these regions for detailed signal profiling. Signal intensities were normalized to RPKM (Reads Per Kilobase per Million mapped reads) to account for differences in sequencing depth and region length. Additionally, we quantified the magnitude of H3K27ac reduction by calculating the fold change in H3K27ac occupancy between BKS Ctrl and db/db mice for each of the three categories.

**HiChIP data processing**

HiChIP data were processed using HiC-Pro[15] using default parameters, except for setting the LIGATION_SITE to GATCGATC, GANTGATC, GANTANTC, GATCANTC and generating GENOME_FRAGMENT using digest_genome.py in HiC-Pro utilities. The following parameters were used: -r ^GATCG ^ANTC -o reference_genome_restriction_sites.bed reference_genome.fasta. Paired-end sequencing reads were first trimmed to remove low-quality bases (Phred score < 20) and adaptor sequences using Trimmomatic[8]. The trimmed reads were then mapped to the hg19 or mm10 reference genome using bowtie2[16] with the parameters "--very-sensitive --end-to-end --reorder". Uniquely mapped reads were assigned to restriction fragments. The raw reads were filtered for valid pairs (VI) which included removal of duplicates, dangling ends, self-circles, and pairs with mapping quality less than 30.

To generate contact matrices, the Hi-C files were created using hicprotojuiebox.py utilities in HiC-Pro[15], with 5 kb, 10 kb and 25 kb resolutions. The normalized interaction matrices were then imported into Juicebox[17] for visualization, where Knight-Ruiz (KR) matrix balancing was applied to correct for experimental and technical biases.

Quality control metrics including the percentage of valid pairs, unique mapping rate, PCR duplicate rate were assessed at each processing step and summazied at Table S1.

HiChIP anchors were defined by intersecting the interaction data with H3K27ac ChIP-seq peaks. Only anchors overlapping H3K27ac peaks were retained, ensuring that each anchor corresponded to a putative regulatory element. HiChIP anchors from all the replicate samples were merged and the number of reads per sample that overlapped the universal anchor set were counted. The overlapping anchors were merged and a single anchor and counts matrix was created. Principal Component Analysis (PCA) was performed on this matrix to assess sample relationships and identify major sources of variation in the chromatin interaction profiles.

**Analysis of differential chromatin interaction intensities at JUNB-bound anchors**

JUNB ChIP peaks were first intersected with HiChIP interaction anchors in both the BKS-Ctrl and BKS-db/db conditions using bedtools intersect[12]. Only anchors overlapping a JUNB peak in either condition were retained for downstream analyses.

Interaction intensities (contact frequencies) for each JUNB-bound anchor were quantified using Juicer tools[18] dump command with KR (Knight-Ruiz) normalized matrices at 5 kb resolution. The interaction frequencies were normalized and compared between conditions to identify differential chromatin interactions associated with JUNB binding sites.

**Integrated analysis of cis-regulatory elements and gene expression changes under JUNB overexpression conditions**

The following ChIP-seq sample groups were processed: Human Umbilical Vein Endothelial Cells (HUVECs) cultured under (1) normal glucose with Null adenovirus (NG), (2) high glucose conditions with null adenovirus (HG), and (3) high glucose conditions with *JUNB* overexpression (HG-JUNB$^{OE}$). PCA across samples were performed using the variance stabilized transformed values (vst) and the ggplot2 package for visualization. Differential H3K27ac peak analysis were performed using DiffBind package (https://bioconductor.org/packages/release/bioc/html/DiffBind.html) for NG versus HG, and HG versus HG-*JUNB*$^{OE}$. EdgeR-normalized ChIP-seq counts across samples and all merged peak set regions were converted to $\log_2$(CPM+1) values. Peaks with |log2FoldChange| > 1 and FDR < 0.05 were considered as differential CREs.

In parallel, differential expression analysis was performed between NG versus HG and HG versus HG-*JUNB*$^{OE}$ groups to identify differentially expressed genes (DEGs). Genes with |log2FoldChange| > 1 and adjusted p-value < 0.05 were considered as differentially expressed genes (DEGs).

A Sankey diagram was generated to visualize genes and CREs that were downregulated in HG and subsequently upregulated in HG-*JUNB*$^{OE}$. The identified CREs were functionally annotated using Genomic Regions Enrichment of Annotations Tool (GREAT)(Version 4.0.4)[19] with default "basal plus extension" parameters: 5 kb upstream, 1 kb downstream, and up to 1000 kb for distal elements), applying the hg38 assembly. Enriched biological pathways were identified using a hypergeometric test with FDR correction (q-value < 0.05). Functional annotation analysis was performed using ToppGene Suite[20] with the following parameters: p-value cutoff = 0.05, FDR B&H correction, and minimum gene set size = 10. For visualization, the top 20 most significantly enriched terms in each category were selected and plotted using ggplot2.

### Analysis of RIME (Rapid Immunoprecipitation Mass Spectrometry of Endogenous Proteins) data

The Enriched Proteins List analysis began with processing two technical replicates of experimental samples against an IgG negative control. The data underwent rigorous filtering by removing proteins present in the IgG negative control and those with spectral counts below 5. The analysis generated three comprehensive lists: proteins unique to each of the two replicates and proteins common to both replicates, with spectral counts averaged for the latter. For each identified protein, essential information including gene name, protein name, and spectral count was documented.

### Supplemental Figure Legends

**Figure S1. Characterization of chromatin landscape and JUNB function in diabetic endothelial cells, related to Figure 1.**

A. Principal Component Analysis (PCA) of H3K27ac ChIP-seq profiles from endothelial cells of three diabetic mouse models.

B. Comparison of H3K27ac ChIP-seq signal intensities at JUNB-bound (n = 18,282) versus non-JUNB-bound (n = 22,384) peaks in BKS control (Ctrl) mice.

C-D. GO enrichment analysis of JUNB bound (n = 16,028) and non-JUNB bound (n = 12,613) HiChIP anchors in BSK Ctrl.

E. PCA of H3K27ac HiChIP chromatin loops from endothelial cells of BKS mouse models.

F-G. Genome browser view displaying multi-omics datasets including H3K27ac ChIP-seq signals, H3K27ac HiChIP chromatin interactions, JUNB ChIP-seq signals and RNA-seq expression values at the *Ccnd2* (F) and *Igf1*(G) loci in BKS Ctrl and db/db endothelial cells.

H. Quantification of *JUNB* expression in WT-Ctrl and WT-sh*Junb* by qPCR.

I-J. Representative images for wound healing assay (I), and quantification of wound closure (J) after 24 h in wild-type mouse aortic endothelial cells transduced with control shRNA adenovirus (WT-Ctrl) and transduced with shRNA-*Junb* adenovirus (WT-sh*Junb*). Data were analyzed by Student's *t* test (n = 5).

K-L. Representative images for transwell assay (K) and quantification data of transwell assay (L) in mouse endothelial cells. Data were analyzed by Student's *t* test (n = 5).

M-N. Representative images (M) and quantification data (N) of tube formation assay in mouse endothelial cells. Data were analyzed by Student's *t* test (n = 5).

**Figure S2. JUNB overexpression improves endothelial cell function under high glucose conditions, related to Figure 2.**

A. Genome browser view showing multi-omics datasets in the region chr10:74004190-74129699. Datasets were from NG treated HUVEC cells and human primary endothelial cells.

B. GO enrichment analysis of downregulated genes in NG-*JUNB*$^{KD}$ versus NG-shNull.

C-D. Single cell RNA sequencing reveals the expression of *RASGRP3* (C) and *DDIT4* (D) across multiple cell types from various tissues.

E-F. Western blot analysis (E) and quantitative analysis (F) of JUNB in NG, HG and HG-*JUNB*$^{OE}$ HUVECs. Data were analyzed by One-way ANOVA (n = 3).

G-H. Representative images for wound healing assay (G) and quantification of wound closure (H) after 24 h in NG, HG and HG-*JUNB*$^{OE}$ HUVECs. Data were analyzed by One-way ANOVA (n = 5).

I-J. Representative images for transwell assay (I), and quantification of migrated cell numbers (J) in NG, HG and HG-*JUNB*$^{OE}$ HUVECs. Data were analyzed by One-way ANOVA (n = 5).

K-L. Representative images for tube formation (K), and quantification of junction numbers (L) in NG, HG and HG-*JUNB*$^{OE}$ HUVECs. Data were analyzed by One-way ANOVA (n = 5).

**Figure S3. Analysis of JUNB-interacting proteins and JUNB ChIP-seq data, related to Figure 3.**

A. Quantification of *Junb* expression in Ctrl and db/db BKS by qPCR.

B. Quantification of *JUNB* expression in NG and HG HUVECs by qPCR.

C-D. Venn diagram displaying the overlap of proteins identified for Rapid Immunoprecipitation Mass Spectrometry of Endogenous Proteins (RIME) by anti-JUNB and anti-IgG antibodies in NG (C) and HG (D) HUVECs.

E. Table listing enriched proteins identified by RIME in both NG and HG HUVECs. The enriched proteins were categorized into six functional subtypes by their GO annotation. Ratio in NG represents the relative expression ratio that was calculated by dividing the spectrum counts of enriched protein by that of JUNB in NG HUVECs. Ratio in HG represents the corresponding ratio calculated in HG HUVECs.

**Figure S4. RBBP6 knockdown enhances endothelial cell function under high glucose conditions, related to Figure 4.**

A-B. Single cell RNA sequencing reveals the expression of *MANSC1* (A) and *ZDHHC6* (B) across multiple cell types from various tissues.

C-D. Representative images for transwell assay (C) and quantification of migrated cell number (D) in NG-shNull, HG-shNull and HG-*RBBP6*$^{KD}$ HUVECs. Data were analyzed by One-way ANOVA (n = 5).

E-F. Representative images of wound healing assay (E), and quantification of wound closure (F) after 24 h in NG-shNull, HG-shNull and HG-*RBBP6*$^{KD}$ HUVECs. Data were analyzed using One-way ANOVA (n = 5).

G-H. Representative images for tube formation assay (G) and quantification of junction number (H) in NG-shNull, HG-shNull and HG-*RBBP6*$^{KD}$ HUVECs. Data were analyzed by One-way ANOVA (n = 5).
